# Hydrodynamic instabilities in membrane systems with current loading, Fourier analysis

**DOI:** 10.1101/2025.10.07.681044

**Authors:** Sławomir Grzegorczyn, Paweł Dolibog, Iwona Dylong, Andrzej Ślęzak

## Abstract

The article presents the results of current measurements in membrane systems with bacterial cellulose membranes, located in horizontal plane, for various initial quotients of NaCl concentrations on the membrane. The obtained time-current characteristics indicate a stable formation of concentration boundary layers near the membrane for the configuration with a solution of lower concentration and lower density above the membrane (A). In turn, for the configuration with a higher concentration and higher density solution above the membrane (B) and a sufficiently large initial concentration quotient on the membrane, greater than 50, current pulsations are observed over time, resulting from hydrodynamic instabilities occurring in the vicinity of the membrane. The increase of initial concentration quotient on the membrane in configuration B causes an increase in the frequency of current pulsations and a change in their amplitudes over time. Furthermore, significant differences were observed between the types of the temporal changes in membrane currents in both configurations, and these differences persisted even 24 hours after turning off mechanical stirring of solutions. To analyze the hydrodynamic instabilities in configuration B of the membrane system, Fourier analysis was used both in the range of observed pulsations of the measured currents (from 50 to 250 min) and in the twenty-minute intervals with the intervals centered at 20, 100, 150, 200 and 290 min). As the analysis shows in all tested time ranges, the average signal power of currents in the frequency range from 0.05 to 1 min^-1^, depends non-linearly on the initial concentration quotient on the membrane, showing a maximum for the concentration quotient on the membrane equal to 2500.

## Introduction

Bacterial cellulose membranes (*Biofill*), due to their structure characterized by high porosity and degree of hydration, have good transport properties for both electrolytes and non-electrolyte substances. For this reason, they have found applications in solving various technical [1–3] as well as medical [4,5] problems. Due to their impermeability to bacteria and, at the same time, easy exchange of water and small molecules, thin sheets of bacterial cellulose are used as dressings for hard-to-heal wounds [6]. Bacterial cellulose is an interesting and increasingly popular material due to its significantly smaller cellulose fibers and fewer other components compared to plant cellulose. For this reason, bacterial cellulose has found numerous applications in medicine [7] and technology [8,9]. The membrane itself constitutes a barrier (physical, structural, electrical) to the flow of solutions between the separated compartments, which allows for controlled transport of substances in single- and multi-membrane systems. Due to their flow-limiting properties and the ability to control the fluxes of substances, membranes have found wide practical applications, including seawater desalination [10,11], wastewater treatment in both industrial and municipal processes [12], electricity generation [10,13,14], and others. In each of these applications, membrane concentration polarization is important, and research is currently underway to limit its impact on membrane transport.

One of the important models describing the transport of non-electrolyte substances and electrolytes through membranes is the Kedem-Katchalsky (KK) model [15]. This model assumes homogeneous solutions on both sides of the membrane, most often ensured by mechanical stirring of solutions. The essence of the model is the equations

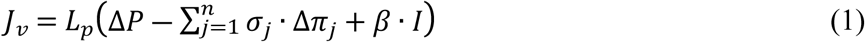

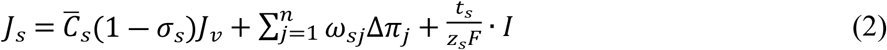

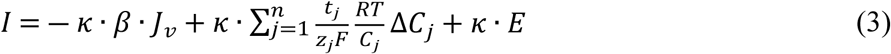

where *J_v_* and *J_s_* are the volume and ion fluxes (*s* – indexes for suitable ions, *n* number of ions in solution), *I* is the density current through the membrane, Δ*P* is the difference of mechanical pressure through the membrane, Δ*π*_*j*_ is the osmotic pressure difference through the membrane for *j* solute, 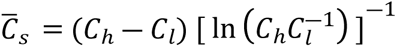 is an average *s* solute concentration in the membrane and 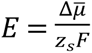 is the gradient of electrical potential on the membrane. Besides, *C*_h_ and *C*_l_ (*C*_h_ > *C*_l_) are the solute concentrations in the chambers at the initial moment, *L*_p_, *σ*_s_ and *ω*_s_ are hydraulic permeability, reflection and solute permeability coefficients for membrane suitably. *β*, *t_s_* and *κ* are electroosmotic coefficient, transference number of ions *s* and conductivity of the membrane suitably. Besides *F*, *R* and *T* are the Faraday number, gas constant and absolute temperature suitably, 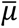 is the electrochemical potential of solution and *z*_*s*_is the valence of ion s.

In practice, the phenomenon of concentration polarization of the membrane occurs, which causes heterogeneity of solutions near the membrane surface. This requires extending the KK model to the case of heterogeneous solutions [16]. One possible extension of the KK model is the model of solutions layers in membrane chambers using the diffusion equation for solutions transport in chambers. The diffusion equation can be shown as

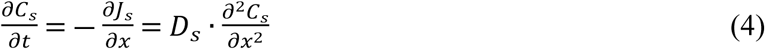

where *D_s_* is the diffusion coefficient of solute *s* in aqueous solutions. As previously mentioned, one of the important factors influencing the transport of substances through the barrier created by the membrane is the phenomenon of concentration polarization of the membrane [16,17]. This phenomenon is associated with the appearance of concentration gradients of transported substances, perpendicular to the membrane surface and significantly decreasing the fluxes of transported substances [17,18]. The associated layers with concentration gradients in the vicinity of membranes, called concentration boundary layers (CBLs) [19,20], are often investigated for potential reduction of concentration polarization of membrane [21,22] and thus decreasing their impact on membrane transport. Another phenomenon that may appear in CBLs are hydrodynamic instabilities [18], which may be caused by solution density gradients directed opposite to the gravitational field strength. Other causes of hydrodynamic instabilities related to density disturbances may be large currents flowing through solutions [23,24] or chemical reactions occurring in solutions [25–27]. Disturbance of the density near the membrane by various factors can lead to the blurring of CBLs and causing a significant reduction in the concentration polarization of the membrane and thus an increase in substances fluxes through the membrane. Density gradients near membrane can be caused by temperature gradients or by concentration gradients of substances whose densities depend on the concentration of the substance. At sufficiently large density gradients, additional convective fluid motions can occur, leading to hydrodynamic instabilities near the membrane [28–30]. A parameter that is crucial in the boundary conditions where hydrodynamic instabilities arise in CBLs is the critical Rayleigh number. The Rayleigh number itself can be represented as [31]

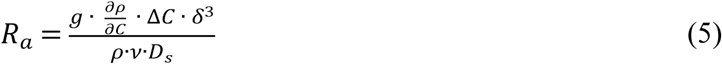

where *g* is the gravitational acceleration, Δ*C* is the difference of concentrations in CBL, *δ* is the thickness of CBL, *ρ* and *ν* are the density and the kinematic viscosity coefficient of solution suitably, *D*_s_ is the diffusion coefficient for solute *s* in solution. Critical value of the Rayleigh number depends on the type of system in which the displacements of substances are considered [32]. The value of the Rayleigh number allows determining the stability conditions of solutions in CBLs and the type of flows observed in the chambers of the membrane system. In the case of membranes, when the membrane surface can be considered as a fixed rigid CBL border, while the second CBL boundary is blurred, the limiting value of the Rayleigh number is taken as 1100.6 [32]. Below this value, the solution flow through the CBL is laminar (diffusive creation of CBLs), whereas after exceeding the critical value of the Rayleigh number, the diffusion flows are disturbed by turbulent displacements of solutions in the CBLs (hydrodynamic instabilities in CBLs).

The method we use to study the dynamics of CBL evolution in membrane systems is a method based on the electrodes that are reversible with respect to the electrolyte (e.g. Ag|AgCl electrodes for chloride solutions) [20,33]. It allows the measurement of voltages between electrodes immersed directly in solutions, using a millivoltmeter with high internal resistance. The voltage between the electrodes in the membrane system, immersed directly in the solutions, can be represented as [33,34]

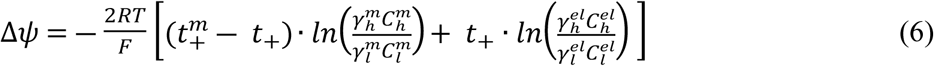

where: 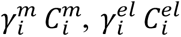 are the products of ion activity coefficients and concentrations suitably at membrane surfaces and electrode surfaces (*h* or *l*), 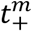 and *t*_+_ are the apparent transference numbers for Na^+^ ions in membrane and solution in chamber suitably, *R*, *T* and *F* are the gas constant, thermodynamic temperature and Faraday constant.

Time characteristics of voltages in membrane systems enable the study of diffusional CBL reconstruction phenomenon and the determination of the conditions of appearance and the type of instability in CBLs [16,28,35]. Measurement of the time characteristics of membrane currents, with the use of an electrical meter with significantly lower resistance should allow for the expansion of the scope of these studies. In this case, electrical measurements in a membrane system would occur with significantly greater charge transport through the membrane and CBLs, i.e., under conditions of significantly higher current load. The measured membrane current can be presented in the form [36]

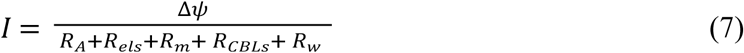

As can be seen from the above equation, the membrane current will depend on the voltage generated at the electrodes immersed directly in the solutions (Δ*ψ*), as well as on the resistance of the membrane itself (*R*_*m*_) and the solutions in CBLs (between electrodes and membrane - *R*_*CBLs*_), as well as the resistances of the electrodes (*R*_*els*_), the connecting wires (*R*_*w*_) and the internal resistance of the nanoammeter (*R*_*A*_). For this reason, the dynamics of voltage changes over time in membrane systems with negligible currents may differ significantly from the dynamics of membrane currents measured under different concentration conditions in the membrane system. This will allow the analysis of transport conditions in membrane systems with large charge transports through the membrane.

In our case, the main direction of interest was focused on hydrodynamic instabilities in CBLs and their characterization by time dependence of currents in membrane systems with membrane in the horizontal plane and mainly in the configuration in which the higher density solution was located above the membrane (configuration B). The stability conditions of solutions in CBLs are determined by the density gradients in solutions (caused by concentration gradients), which influence the Rayleigh number defined by the equation (5). Their nature, reflected in the time characteristics of membrane currents for different initial conditions, should depend on these conditions, i.e. the initial solutes concentrations quotient on both sides of the membrane. Due to the fact that the basic manifestation of hydrodynamic instabilities in the membrane system are pulsations of the measured electrical parameters [37,38], we decided to use the method of analysis of the time-varying current signal - the Fast Fourier Transform (FFT). The analysis was carried out for time intervals in which hydrodynamic instabilities appear, visible as pulsations of the measured currents. The results of the analysis were presented using the appropriate transformation spectra of these time intervals. To visualize the Fourier analysis of the observed processes for different initial conditions, the average signal power of the Fourier spectrum was determined in the frequency range in which the signal powers were at least an order of magnitude higher than in the remaining frequency range. The duration of individual pulsations observed under different measurement conditions also provided a suggestion for selection of this range.

The aim of this paper is to expand the description of the dynamics of transport processes in the membrane systems with large currents and to characterize hydrodynamic instabilities in CBLs visible as time-dependent pulsations of measured currents using FFT.

## Materials and methods

A two-chamber system with a bacterial cellulose membrane (*Biofill* membrane, Fibrocel Productos Biotechnologicos Ltd. Curitiba, Brazil) and a mechanical solution stirring system were used to measure currents in the membrane system. The membrane systems consisted of two cylindrical chambers with volumes 200 cm^3^ each, while the surface of the membrane separating chambers was S = 6.1cm^2^. The transport parameters of the bacterial cellulose membrane for NaCl solutions, determined experimentally [38], were: *L*_p_ = 0.5·10^-11^ m^3^N^-1^s^-1^, *σ*_s_ = 0.06 and *ω*_s_ = 14.3·10^-10^ molN^-1^s^-1^. A nanoammeter with an internal resistance of 0.9 MΩ, a range of 100 nA, and a resolution of 0.1 nA (Meratronik U276) was used and was connected to a computer with software for recording temporal changes in membrane currents. Ag|AgCl electrodes were used (made by repeatedly immersing purified silver wire in molten AgCl and then thermally stabilized). The length of prepared electrodes were 14.0 mm, diameter 1.0 mm, while their electrical resistance depended only slightly on the solution concentration (*R*_*els*_= 0.95MΩ). In turn, the membrane resistance, calculated for example for a NaCl concentration of 0.01 mol m^-3^, is about 5.2 kΩ [36] and is negligibly small compared to the resistance of the electrodes and the internal resistance of the nanoammeter. The entire measurement system was placed in a grounded and thermally stabilized metal enclosure, protecting it from the influence of external electric and electromagnetic fields. The scheme of the measurement system is shown in Fig. 1, in two configurations, A and B, respectively. Configuration A contained the higher density solution below the membrane, while in configuration B the higher density solution was above the membrane.

**Fig. 1.** Scheme of the measuring system with a bacterial cellulose membrane (M) in configurations A and B, with aqueous NaCl solutions with initial concentrations *C*_h_ > *C*_l_, S is the motor of the solution stirring system, nA is the nanoammeter and Comp is a computer.

During measurements of currents, the measuring system with the membrane in the horizontal plane, had two configurations: with a higher-density solution below the membrane (configuration A) and with a higher-density solution above the membrane (configuration B). As a membrane a bacterial cellulose membrane (*Biofill*) was used, and aqueous NaCl solutions of various concentrations were used as the electrolyte. For NaCl solutions, the higher the NaCl concentration, the higher the solution density. This allows to change the conditions for CBL formation over time by selecting appropriate solution concentrations at the initial moment and thus control the conditions for the development of hydrodynamic instabilities in the system. The initial lower concentration in one chamber was set at 10^-5^ mol/l NaCl, while in the second chamber the concentration of the NaCl aqueous solution at the beginning of experiment was varied from 5·10^-5^ mol/l to 7.5·10^-2^ mol/l. To ensure homogeneity of the solutions, the solutions in the chambers were stirred before the measurement (for a maximum of 2 minutes). The moment of turning off the stirring of the solutions in chambers of the membrane system was the starting point for the measurement of membrane current. The current values were measured every 4 seconds for 6 hours, and then, to determine the steady state of the membrane system, the current was additionally measured 24 hours after the turning off mechanical stirring of the solutions. The error in the preparation of NaCl solution concentrations did not exceed 2%. During measurements the entire membrane system was located in a metal, grounded measurement chamber, thermally isolated from the surrounding environment.

The current in the membrane system as a function of time obtained for configuration B, characterized by current pulsations related to hydrodynamic instabilities, were analyzed using the Fourier method (Fast Fourier Transform - FFT). The Origin Pro 2024 was used for Fourier analysis and graphical presentation of obtained results. The time intervals: from 50 to 250 minutes, and smaller 20-minute intervals with the interval centers at 20, 100, 150, 200 and 290 minutes, in which voltage pulsations occurred were analyzed. These analyses were carried out for the membrane system under different initial conditions (different *C*_h_/*C*_l_) and for configuration B, in which hydrodynamic instabilities appeared. Moreover, due to the fact that the main frequencies with non-zero signal power occurred in the frequency range up to 1 min^-1^, a parameter characterizing the average signal power in the range from 0.05 to 1 min^-1^ was also calculated according to the equation

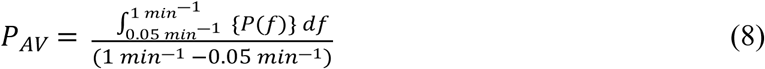

where: *P* - signal power in the frequency domain, f - signal frequency, *P*_AV_ - average signal power in the selected frequency range. The lower integration range was adopted due to the high amplitude of the signal power for low frequency values associated with the continuous decrease in current value over time caused by the gradual disappearance of the concentration forces on the membrane and in the CBLs.

## Results and discussion

Figure 2 presents the time characteristics of membrane currents at different values of the quotients of concentrations on the membrane at the moment of switching off the mechanical stirring of solutions, for the system with a bacterial cellulose membrane. As for the nature of the changes, the curves illustrating the time characteristics of the membrane currents, presented in Figure 3, are similar to the curves illustrating the time characteristics of the voltages between electrodes immersed into solutions, which indicates a significant relationship between the voltage between the electrodes, resulting from location of electrodes in solutions with different NaCl concentrations, and the current flowing through the membrane. The difference between the time characteristics of membrane voltages and currents may be related to different values of the current density flowing through the membrane during measurements and changes in resistance in the membrane system due to changes in the concentrations of solutions in the membrane and CBLs during measurements. When measuring the time characteristics of voltages between electrodes in a membrane system with a millivoltmeter, the maximum current was lower than 1.5 nA, while when measuring the time characteristics of currents in a membrane system with a nanoammeter, the currents flowing were several dozen times greater, but did not exceed 100 nA. This may cause additional concentration disturbances in the membrane and CBLs at sufficiently high current intensities.

**Fig. 2.** Time characteristics of membrane currents (*I*) for the *Biofill* membrane and electrodes at a distance of 5 mm from the membrane surface each, for configurations A (curve A) and B (curve B), for initial NaCl quotients on the membrane (*C*_h_/*C*_l_) equal to: 100 (a), 500 (b), 1000 (c), 2500 (d), 5000 (e) and 7500 (f).

**Fig. 3.** Membrane current intensity as a function of *C_h_/C_l_* on the membrane for bacterial cellulose membranes: *I*_o_ measured at the initial moment during mechanical stirring of solutions, *I*_A_ and *I*_B_ – measured in steady states (24 hours after turning off mechanical stirring of solutions), for configurations A and B, respectively.

Turning off the mechanical stirring of solutions causes the intensity of membrane currents to decrease over time, which is associated with the evolution in time of CBLs. The greatest changes in current intensity are observed in the first few minutes after turning off mechanical stirring of solutions in the membrane system, after which the rate of change in current becomes slower. This results from concentration changes in the membrane and CBLs as a result of ion diffusion through and near the membrane. When the initial concentration ratio on the membrane satisfies the condition *C*_h_/*C*_l_ ≤ 100, the time characteristics of the membrane current after turning off mechanical stirring of solutions are similar in both membrane system configurations. In turn, the time characteristics of membrane currents for configurations A and B and *C_h_/C_l_* ≥ 100 differ significantly. A characteristic feature of the temporal changes in the currents observed in configuration A is the gradual decrease of current over time after turning off the mechanical stirring, which is the effect of diffusional reconstruction of CBLs. For configuration B and for *C_h_/C_l_* ≥ 100, current pulsations appear due to hydrodynamic instabilities in CBLs. The time of observation of current pulsation is longer than 300 minutes from the moment of turning off the mechanical stirring of solutions, with the current pulsation amplitude decreasing systematically during this time. For *C_h_/C_l_* > 1000, current pulsations are characterized by a short pulsation period of current intensity and an initially large amplitude, decreasing over time, which can be associated with the gradual disappearance of thermodynamic forces supporting hydrodynamic instabilities (concentration gradients in the direction of the gravitational field intensity). The short period of current pulsation (approximately 2-3 minutes) indicates quick liquid displacement processes in the CBLs (and near the electrode surfaces), resulting from the emerging density gradients directed opposite to the direction of the gravitational field intensity. These processes, leading to fast stirring of layers of different densities, contribute to the disappearance of the thermodynamic forces (the difference in density between adjacent layers), and thus to a reduction in the amount of mass exchanged between the layers. This results in smaller concentration changes at the electrode surfaces and, consequently, a reduction in the amplitude of pulsations of currents flowing through the membrane.

Figure 3 presents the concentration characteristics of membrane currents for bacterial cellulose membranes, measured in steady state 24 hours after turning off the mechanical stirring of solutions. The current intensities in the membrane system during mechanical stirring of solutions are designated as *I*_o_. In turn, the currents measured 24 hours after turning off the mechanical stirring were marked for configuration A by *I*_A_ and for configuration B by *I*_B_. The current values for *I*_o_ correspond to the average of currents for both configurations of the membrane system during mechanical stirring of solutions.

Figure 3 shows that the increase in *C_h_/C_l_*at the initial moment is the reason for the increase in the current flowing through the membrane during mechanical stirring of solutions in the chambers, to a value of about 70 nA for *C_h_/C_l_* = 7500. Higher *C_h_/C_l_*values at the initial moment mean higher concentration gradients on the membrane at the beginning and appearing over time in the CBLs, which is associated with higher ion fluxes diffusing through the membrane and through the CBLs. These factors affect the higher observed current intensities in the membrane system. Moreover, for *C_h_/C_l_* lower than 25 (*C_h_* = 2.5·10^-4^ mol/l) there are no significant differences between the steady-state membrane currents for configurations A and B. Increasing *C_h_/C_l_* during mechanical stirring of solutions at initial moment above 25 causes the steady-state currents for configuration A to be lower than in B. As it results from the analysis of the time-dependent characteristics of currents in the membrane systems, current pulsations are observed for *C_h_/C_l_* greater than or equal to 100. However, differences in the steady-state currents appear above 25. The reason for the differentiation is the hydrodynamic instabilities caused by gravity, however, at smaller thermodynamic forces in CBLs they are not observed in the form of current pulsations, but are seen in steady states for both configurations of the membrane system. In both configurations, CBLs are reconstructed by diffusion, while hydrodynamic instabilities occurring in configuration B cause the CBLs to “blur” and are responsible for a slower increase in CBL thickness than in configuration A. Additionally, during current measurements, where the currents are several dozen times higher than during measurements of voltage between electrodes, the flowing currents may cause additional concentration disturbances in the membrane system and ohmic potential drops on the membrane and CBLs. An increase in *C_h_/C_l_*across the membrane at the initial moment increases the difference between the steady-state currents. The maximum difference is observed for a *C_h_/C_l_* of approximately 250. A further increase in *C_h_/C_l_* causes a decrease in the differences between the steady states in configurations A and B. This indicates a decrease in the intensity of the processes generating hydrodynamic instabilities in the areas adjacent to the membrane, which determine the thickness of the CBLs.

To analyze the current pulsations over time caused by hydrodynamic instabilities in the CBLs, a Fast Fourier Transform (FFT) analysis was used. The signal in the range of observed current pulsations (ranging from 50 to 250 min) was analyzed, and in whole time range of current measurement, 20-minute subranges were chosen, which were also analyzed using FFT. The results of the FFT analysis of these ranges are shown in Figures 4 and 5.

**Fig. 4.** Dependence of the signal power (*P*) on the frequency, obtained as a result of FFT, for configuration B of the membrane system and time interval 50-250 min, for *C*_h_/*C*_l_ equal to: 100 (a), 500 (b), 1000 (c), 2500 (d), 5000 (e) and 7500 (f), respectively.

**Fig. 5.** Dependence of the signal power (*P*) on the frequency, obtained as a result of FFT, for configuration B of the membrane system and exemplary time interval 90-110 min, respectively for *C*_h_/*C*_l_ equal to: 100 (a), 500 (b), 1000 (c), 2500 (d), 5000 (e) and 7500 (f).

As can be seen from graphs 4 a-f and 5 a-f, the main component of the signal power is in the low frequency range up to 1 min^-1^. This is consistent with the graphs presented in Figure 3 (for configuration B), which show that the current pulsation periods are usually in the range of few to several dozen minutes. The main feature of the FFT spectra, both for the 50-250 minute range (Fig. 4) and for the 20 minute ranges (example graphs in Fig. 4), is that with the increase in the initial NaCl concentration ratio on the membrane (*C*_h_/*C*_l_), the power values initially increase and above *C*_h_/*C*_l_ = 2500 they decrease. Moreover, the frequency range with high power values also shifts towards higher frequencies and above *C*_h_/*C*_l_ = 5000 towards lower ones. The main frequency ranges for which the signal strengths were significantly higher than the others were within the frequency ranges up to 0.6 min^-1^ for the longer interval and from 0.8 to 1 min^-1^ for the narrower (20-minute) intervals. For this reason, in order to capture and better present these relationships, the average power in the range of observed maximum signal powers, i.e. from 0.05 to 1 min^-1^, was defined as the parameter. The lower range was assumed due to the longest observed periods of current pulsation in the time characteristics of currents in the membrane system, and the upper range was assumed as a cut-off from the frequency range with negligible signal powers and no observation of pulsations with frequencies greater than 1 min^-1^ (period shorter than one minute). The signal power averaging range corresponds to the borders of pulsation periods: 20 min and 1 min, respectively. For this interval, the average power in the interval was calculated based on the formula (8) as the integral of the signal power in the selected interval, divided by the interval width. Moreover, the averaging limit at low frequencies was chosen due to the large signal component at very low frequencies resulting from the gradual decrease in the membrane current value over time. Figure 6 shows the dependence of the average power in the frequency range from 0.05 to 1 min^-1^ in selected time-intervals on the initial *C*_h_/*C*_l_ conditions for the bacterial cellulose membrane and aqueous NaCl solutions.

**Fig. 6.** Average power in the frequency range from 0.05 to 1 min^-1^ as a function of *C*_h_/*C*_l_ obtained for FFT and for configuration B of the membrane system and NaCl aqueous solutions. Time intervals of the signal *I*(*t*): from 50 to 250 min (a) and 20-minute intervals (b) with the time interval centered at: 20min (1), 100min (2), 150min (3), 200min (4) and 290min (5)

As can be seen from Figure 7, for both large and small time intervals, the average signal power shows a maximum for *C*_h_/*C*_l_ = 2500. Moreover, Figure 7b shows that as the center of the 20-minute interval is moved towards a longer time, the average signal power in the interval decreases, while in ranges 100 and 150 min. average powers are similar. This can be related with similar intensities of hydrodynamic instabilities in the middle time ranges (100 and 150 min.) of observed pulsations and then a gradual reduction of intensity of hydrodynamic instabilities. The CBLs formation processes are influenced by two main processes: solute transport through the membrane and solute diffusion through the CBLs. These processes lead to an increase in the CBLs thickness over time and possibility of appearance of hydrodynamic instability in the CBLs, observed only in configuration B, which causes a gradual blurring of CBLs outer border. The first process leads to a gradual decrease in the current flowing in the membrane system while the second leads to the appearance of pulsations in time characteristics of measured currents. Because we focus on the analysis of hydrodynamic instabilities, in order to eliminate the systematic decrease of membrane currents resulting from the first process, the differential method was used. This method consists in replacing the points of the measurement series with the differences between the points of the time series that are separated in time by the same interval. We used the algorithm: Δ*I*(*t*_i_) = *I*(*t*_i_) – *I*(*t*_i_+1min.). The time interval between the points of amounts one minute because no pulsations with a period of less than 1 minute are observed. For smaller values of the time interval, the graphs are similar but with a smaller amplitude and a larger number of high-frequency components not observed in the current signal. The graphs obtained in this way for different initial concentrations quotients (*C*_h_/*C*_l_) are presented in Figure 7.

**Fig. 7.** Time characteristics of differences between elements of the series of current intensities (Δ*I*) for configuration B of the membrane system and for the initial NaCl concentrations quotients *C*_h_/*C*_l_ on the membrane: 100 (a), 500 (b), 1000 (c), 2500 (d), 5000 (e) and 7500 (f).

As can be seen from Figure 7, the pulsations in the graphs correspond to the pulsations of the time dependencies of the currents in configuration B shown in Figure 2. Figure 7 shows that the decreasing trend of the current intensity over time has been eliminated, which was the purpose of used algorithm. The lack of a trend line in time characteristics should reduce very low-frequency components in the FFT spectrum. Figures 8 (analysis of the time range from 50 to 250 minutes) and 9 (analysis of exemplary 20-minute time interval: from 90 to 110 min) present graphs of the FFT analysis of the time characteristics of the series of current differences for the initial quotients of NaCl concentrations with the corresponding *C*_h_/*C*_l_ equal to 100 (a), 500 (b), 1000 (c), 2500 (d), 5000 (e) and 7500 (f) respectively.

**Fig. 8.** The dependence of the signal power (*P*) on the frequency, obtained as a result of FFT for configuration B of the membrane system and for time interval 50-250 min., for *C*_h_/*C*_l_ equal to: 100 (a), 500 (b), 1000 (c), 2500 (d), 5000 (e) and 7500 (f), respectively.

Figures 8 and 9 show that there are frequency bands with higher signal power values than the powers in the lower and higher frequency ranges. Increasing *C*_h_/*C*_l_ causes an increase in power amplitudes and above *C*_h_/*C*_l_ = 2500 a decrease in power is observed. Besides, a shift of this band towards higher frequencies can also be observed with increase of *C*_h_/*C*_l_. For the frequency characteristics shown in Figures 8 and 9, the average powers were calculated according to the equation (8) in the frequency range from 0.05 to 1 min^-1^. The dependencies of calculated averaged powers on *C*_h_/*C*_l_ are presented in Figures 10a (for time range from 50 to 250min.) and 10b (for 20-minutes intervals), respectively.

**Fig. 9.** The dependence of the signal power (*P*) on the frequency, obtained as a result of FFT, for configuration B of the membrane system and exemplary time interval 90-110 min, for *C*_h_/*C*_l_ equal to: 100 (a), 500 (b), 1000 (c), 2500 (d), 5000 (e) and 7500 (f), respectively.

**Fig. 10.** Average power in the frequency range from 0.05 to 1 min^-1^ as a function of *C*_h_/*C*_l_ for configuration B of the membrane system and NaCl aqueous solutions, obtained as a result of FFT in time intervals of the signal Δ*I*(*t*): from 50 to 250 min (a) and in 20-minute intervals (b) with the center of the time intervals at: 20min (1), 100min (2), 150min (3), 200min (4) and 290min (5).

As results from Figures 10a and 10b, the maximum average signal power occurs for *C*_h_/*C*_l_=2500 for all analyzed by FFT time intervals. Shifting of the center of smaller time interval to higher values causes that average power of signals in 20-ty minutes intervals (10b) changes over the entire range of the analyzed concentration ratios at the initial moment, with a tendency to decrease the average signal power.

## Conclusions

- The time characteristics of membrane currents in configuration A of the membrane system indicate stable, diffusive reconstruction of CBLs, regardless of the initial quotient of NaCl concentration on the membrane (*C*_h_/*C*_l_). Such characteristics were also observed for configuration B and an initial quotient of NaCl concentration on the membrane lower than 100.
- For configuration B and *C*_h_/*C*_l_ ≥ 100, pulsations of membrane currents were observed, which indicated the appearance of hydrodynamic instabilities connected with density gradients in CBLs
- In steady states of current intensities, measured 24 hours after turning off the mechanical stirring of solutions, differences in currents were observed between two configurations of the membrane systems, with higher currents observed in configuration B. The largest current differences in steady states for configurations A and B were observed in the range of *C*_h_/*C*_l_: from 100 to 500.
- Fourier analysis showed that the average signal power in the frequency range observed in the time graphs increases with increasing *C*_h_/*C*_l_, reaching maximum for *C*_h_/*C*_l_ = 2500 regardless of the analyzed time range.
- For configuration B and the fixed *C*_h_/*C*_l_ the current pulsations do not change or increase slightly with time and for *t* > 150 min. from turning off the mechanical stirring of solutions they decrease with time, which is also confirmed by the FFT analysis in short-time intervals.
- Application of the difference method to membrane currents allows for the removal of the decreasing trend associated with the diffusional reconstruction of CBLs, allowing for a more effective Fourier analysis of hydrodynamic instabilities in CBLs manifested by current pulsations over time.

## Supporting information

**S1-2 Fig 1.** Scheme of the measuring system with a bacterial cellulose membrane (M) in configurations A and B, with aqueous NaCl solutions with initial concentrations *C*_h_ > *C*_l_, S is the motor of the solution stirring system, nA is the nanoammeter and Comp is a computer.

**S3-8 Fig 2.** Time characteristics of membrane currents (*I*) for the *Biofill* membrane and electrodes at a distance of 5 mm from the membrane surface each, for configurations A (curve A) and B (curve B), for initial NaCl quotients on the membrane (*C*_h_/*C*_l_) equal to: 100 (a), 500 (b), 1000 (c), 2500 (d), 5000 (e) and 7500 (f).

**S9 Fig 3.** Membrane current intensity as a function of *C_h_/C_l_* on the membrane for bacterial cellulose membranes: *I*_o_ measured at the initial moment during mechanical stirring of solutions, *I*_A_ and *I*_B_ - in steady states (24 hours after turning off mechanical stirring of solutions), for configurations A and B, respectively.

**S10-15 Fig 4.** Dependence of the signal power (*P*) obtained as a result of FFT on the frequency, for configuration B of the membrane system and time interval 50-250 min, for *C*_h_/*C*_l_ equal to: 100 (a), 500 (b), 1000 (c), 2500 (d), 5000 (e) and 7500 (f), respectively.

**S16-21 Fig 5.** Dependence of the signal power (*P*) on the frequency obtained as a result of FFT, for configuration B of the membrane system and exemplary time interval 90-110 min, respectively for *C*_h_/*C*_l_ equal to: 100 (a), 500 (b), 1000 (c), 2500 (d), 5000 (e) and 7500 (f).

**S22-23 Fig 6.** Average power in the frequency range from 0.05 to 1 min^-1^ as a function of *C*_h_/*C*_l_ obtained for FFT and for configuration B of the membrane system and NaCl aqueous solutions. Time intervals of the signal *I*(*t*): from 50 to 250 min (a) and 20-minute intervals (b) with the time interval centered at: 20min (1), 100min (2), 150min (3), 200min (4) and 290min (5)

**S24-29 Fig 7.** Time characteristics of differences between elements of the series of current intensities (Δ*I*) for configuration B of the membrane system and for the initial NaCl concentrations quotients *C*_h_/*C*_l_ on the membrane: 100 (a), 500 (b), 1000 (c), 2500 (d), 5000 (e) and 7500 (f).

**S30-35 Fig 8.** The dependence of the signal power (*P*) on the frequency obtained as a result of FFT for configuration B of the membrane system and for time interval 50-250 min., for *C*_h_/*C*_l_ equal to: 100 (a), 500 (b), 1000 (c), 2500 (d), 5000 (e) and 7500 (f), respectively.

**S36-41 Fig 9.** The dependence of the signal power (*P*) on the frequency obtained as a result of FFT, for configuration B of the membrane system and exemplary time interval 90-110 min, for *C*_h_/*C*_l_ equal to: 100 (a), 500 (b), 1000 (c), 2500 (d), 5000 (e) and 7500 (f), respectively.

**S42-43 Fig 10.** Average power in the frequency range from 0.05 to 1 min^-1^ as a function of *C*_h_/*C*_l_ for configuration B of the membrane system and NaCl aqueous solutions, obtained as a result of FFT in time intervals of the signal Δ*I*(*t*): from 50 to 250 min (a) and in 20-minute intervals (b) with the center of the time intervals at: 20min (1), 100min (2), 150min (3), 200min (4) and 290min (5).

**S1 File** Raw data for article.xlsx

---

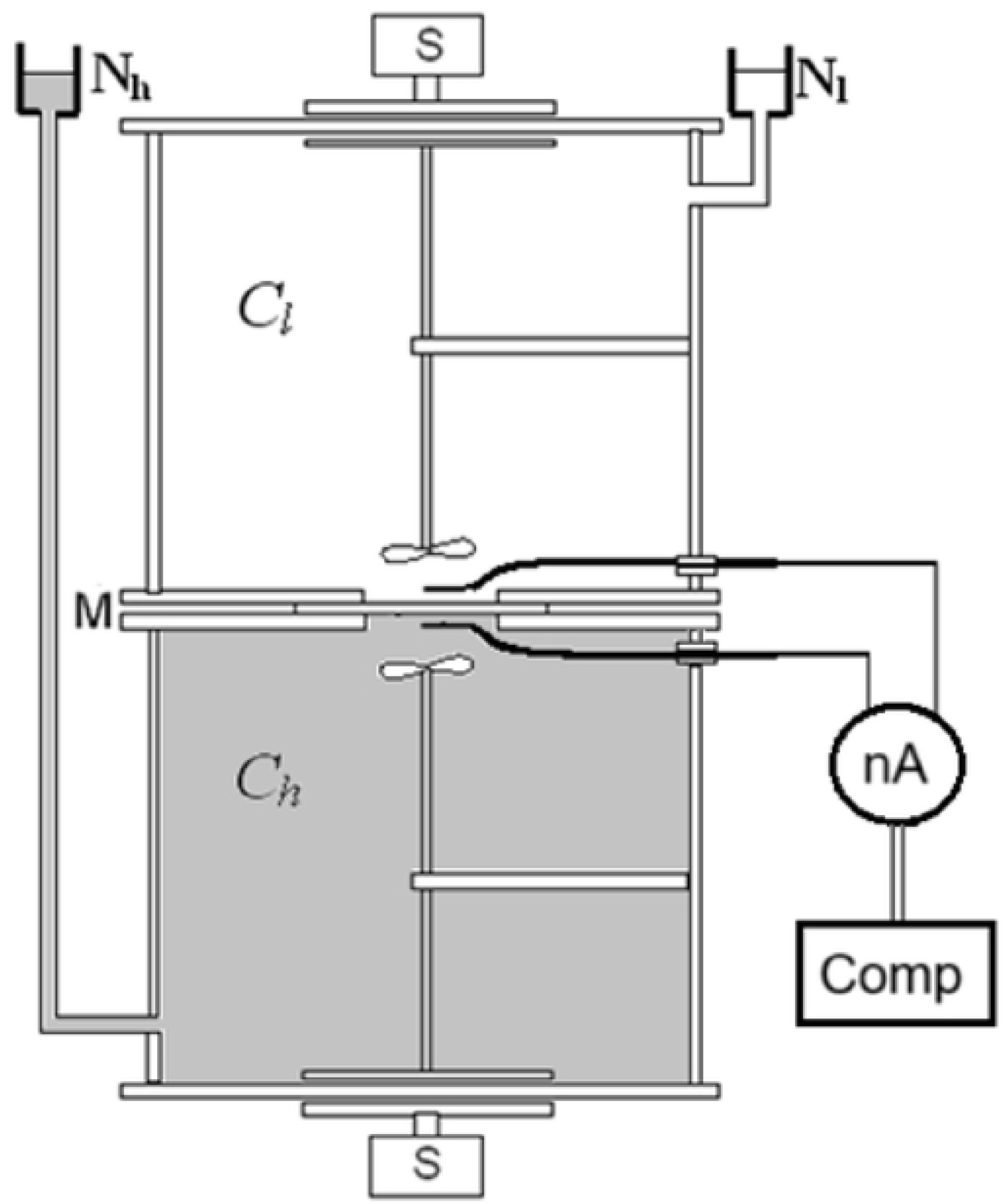

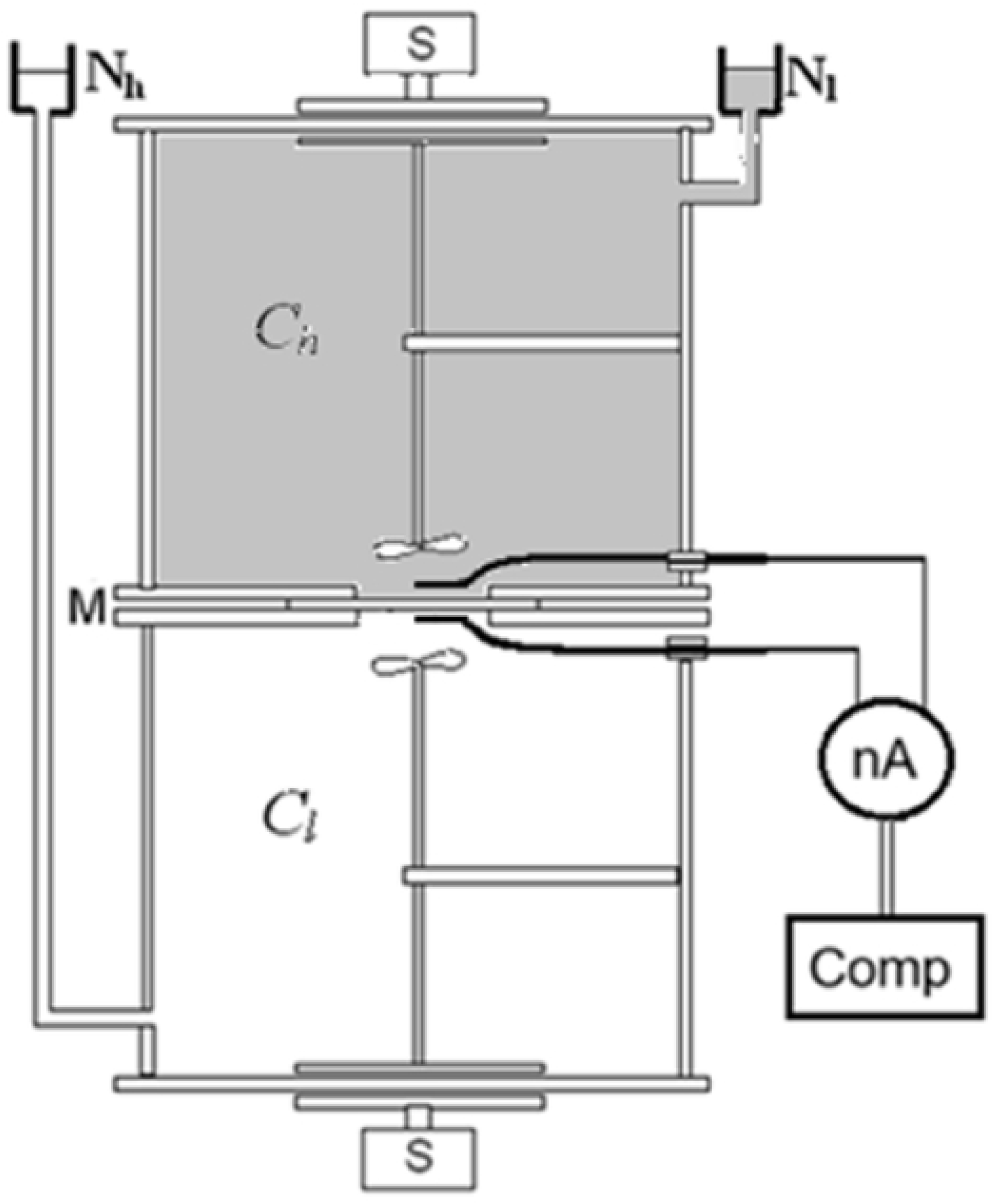

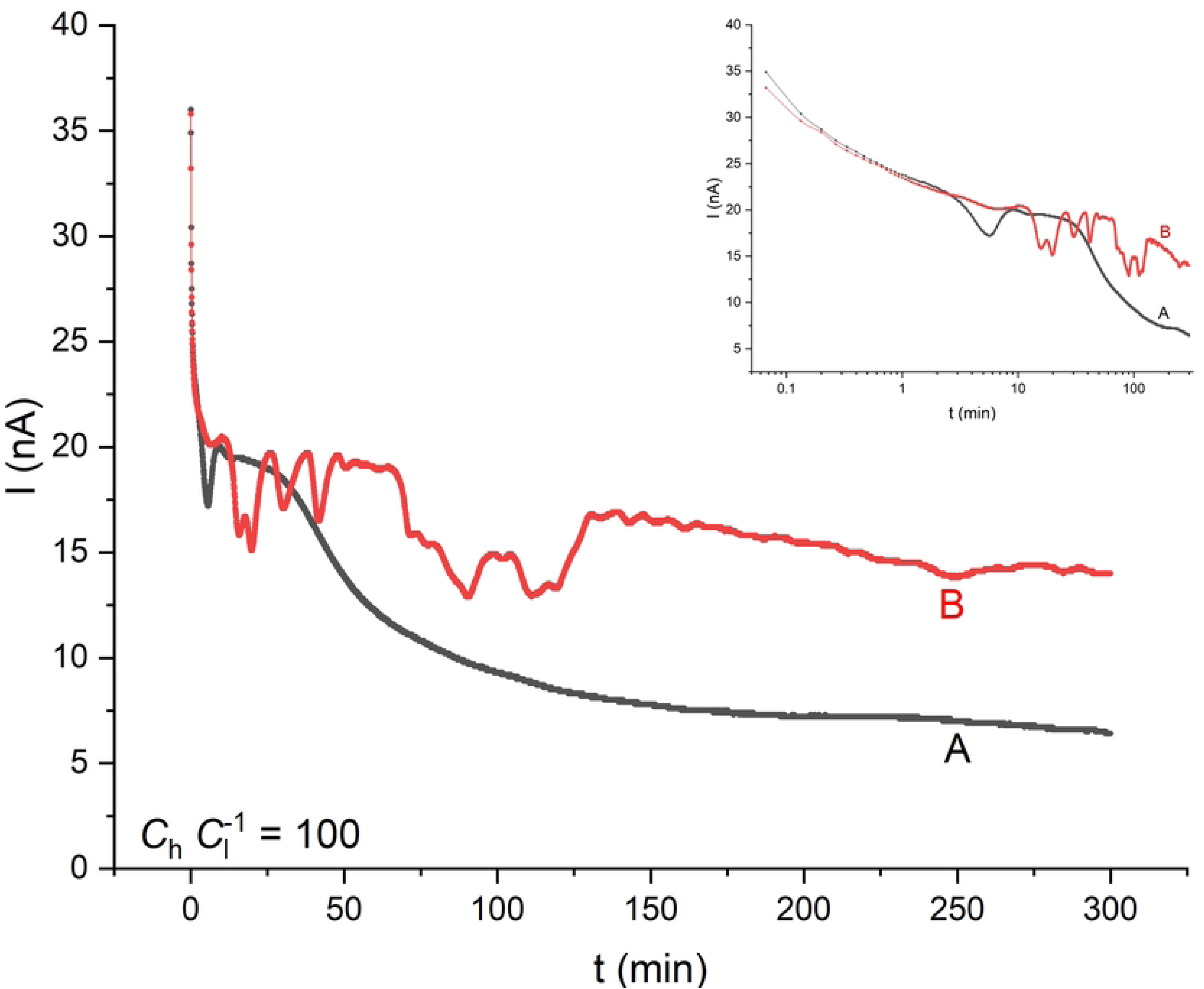

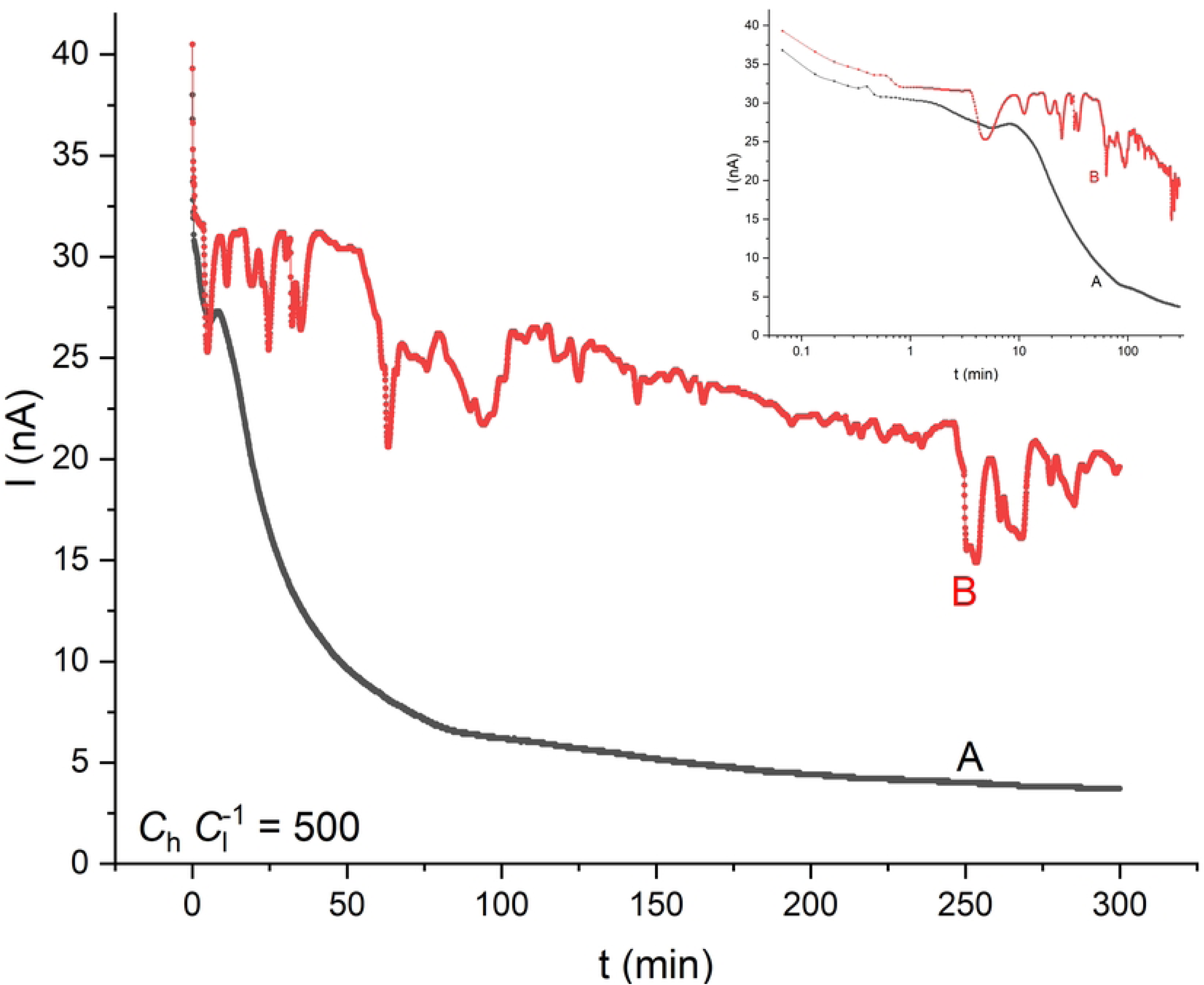

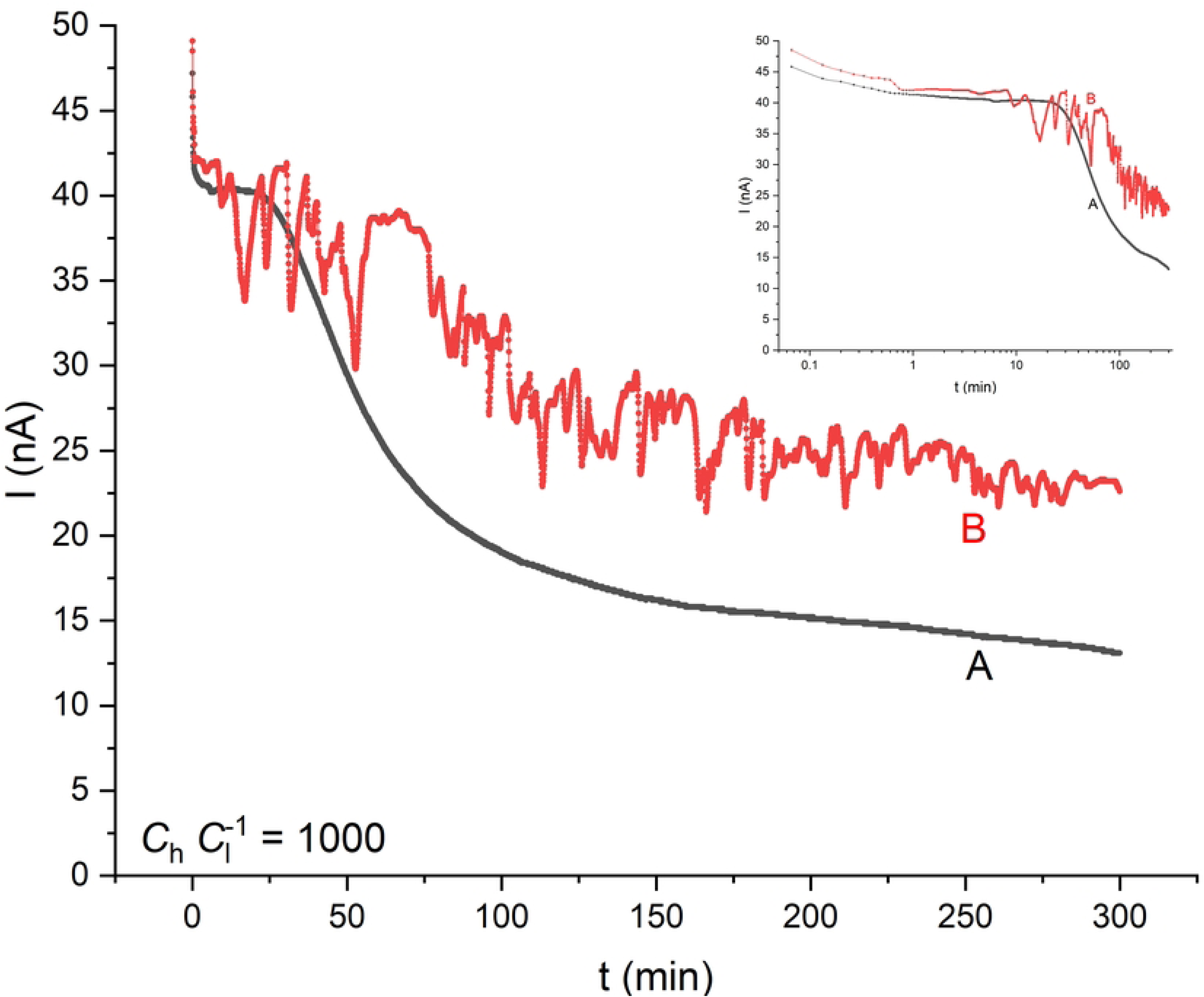

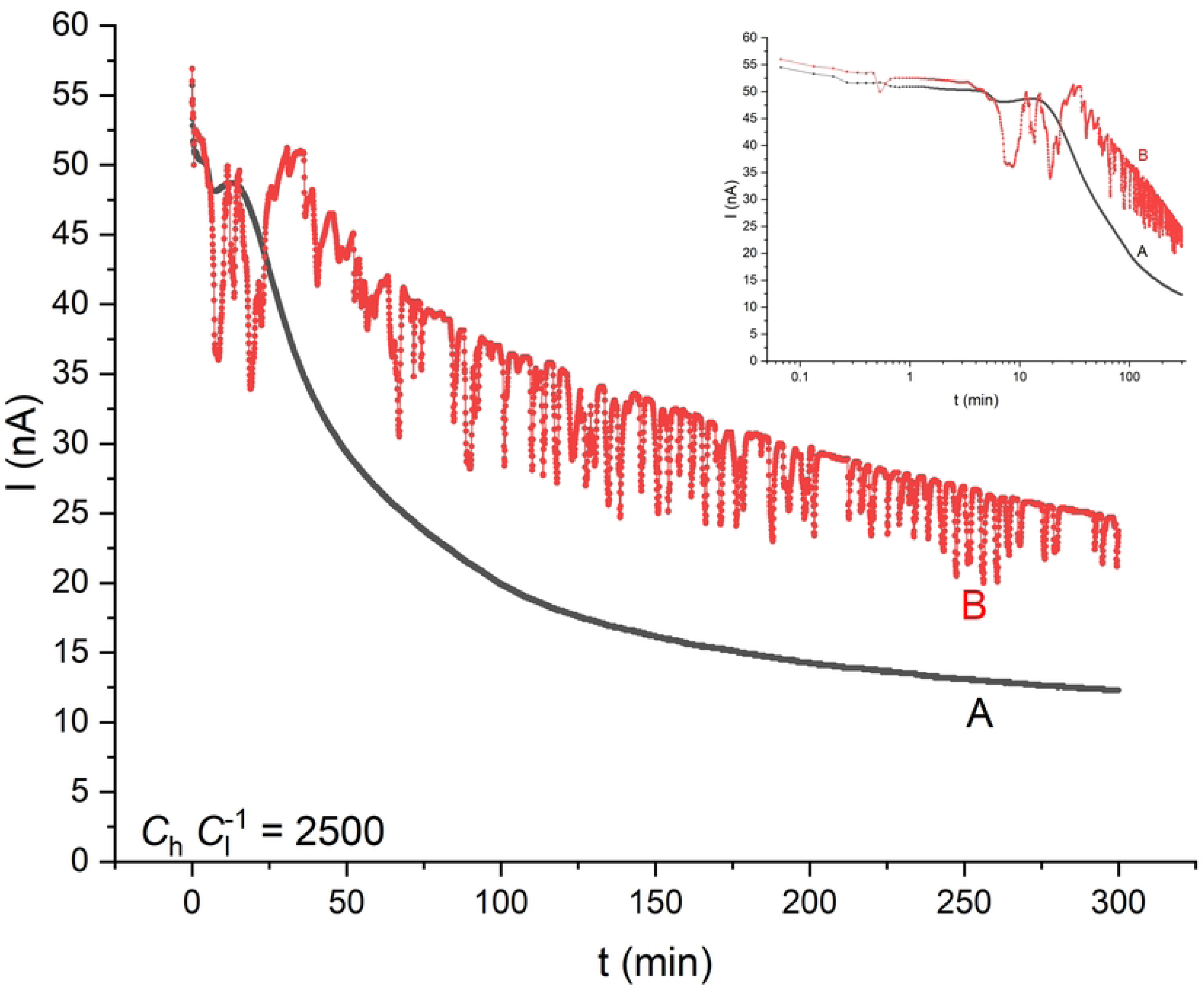

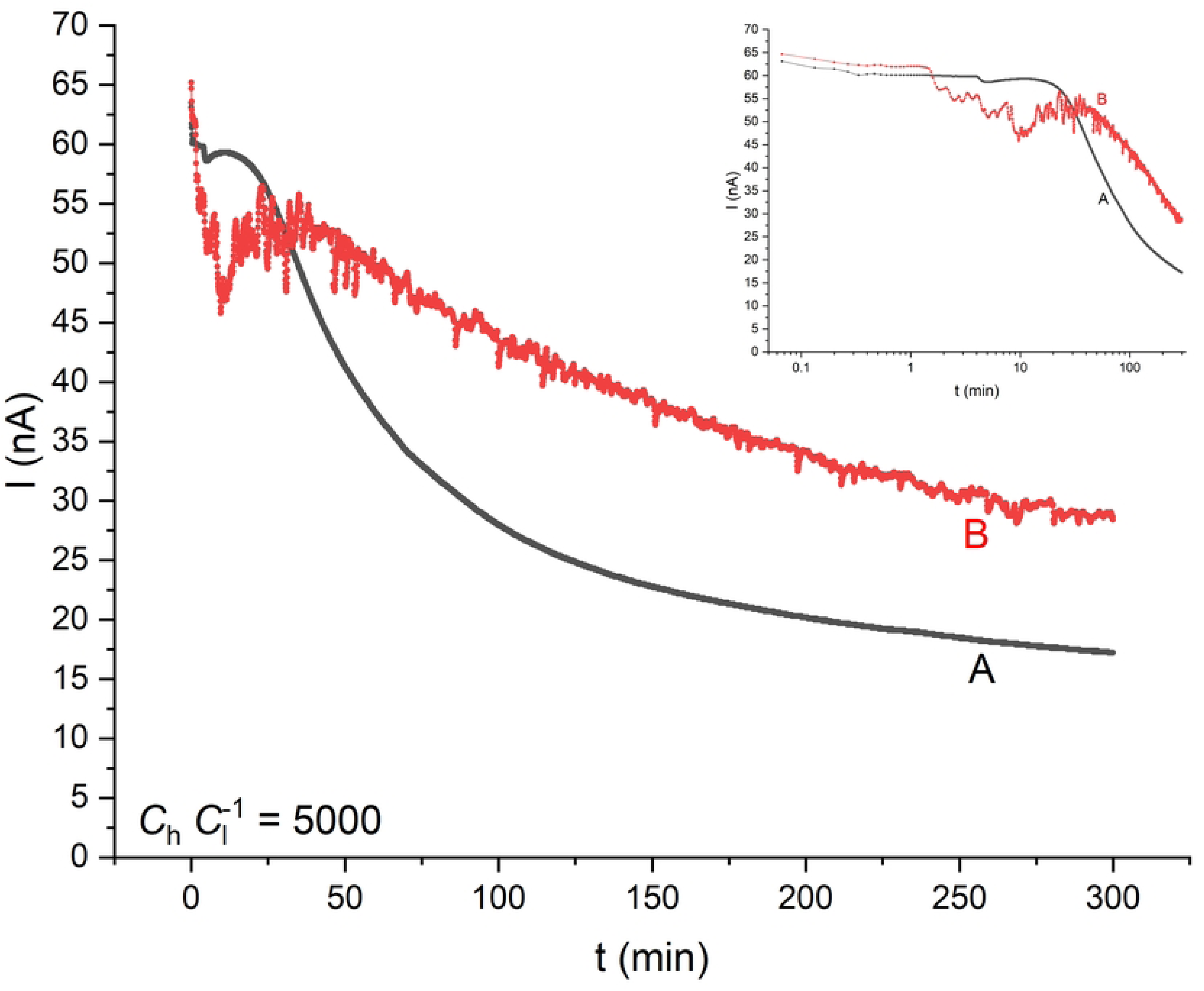

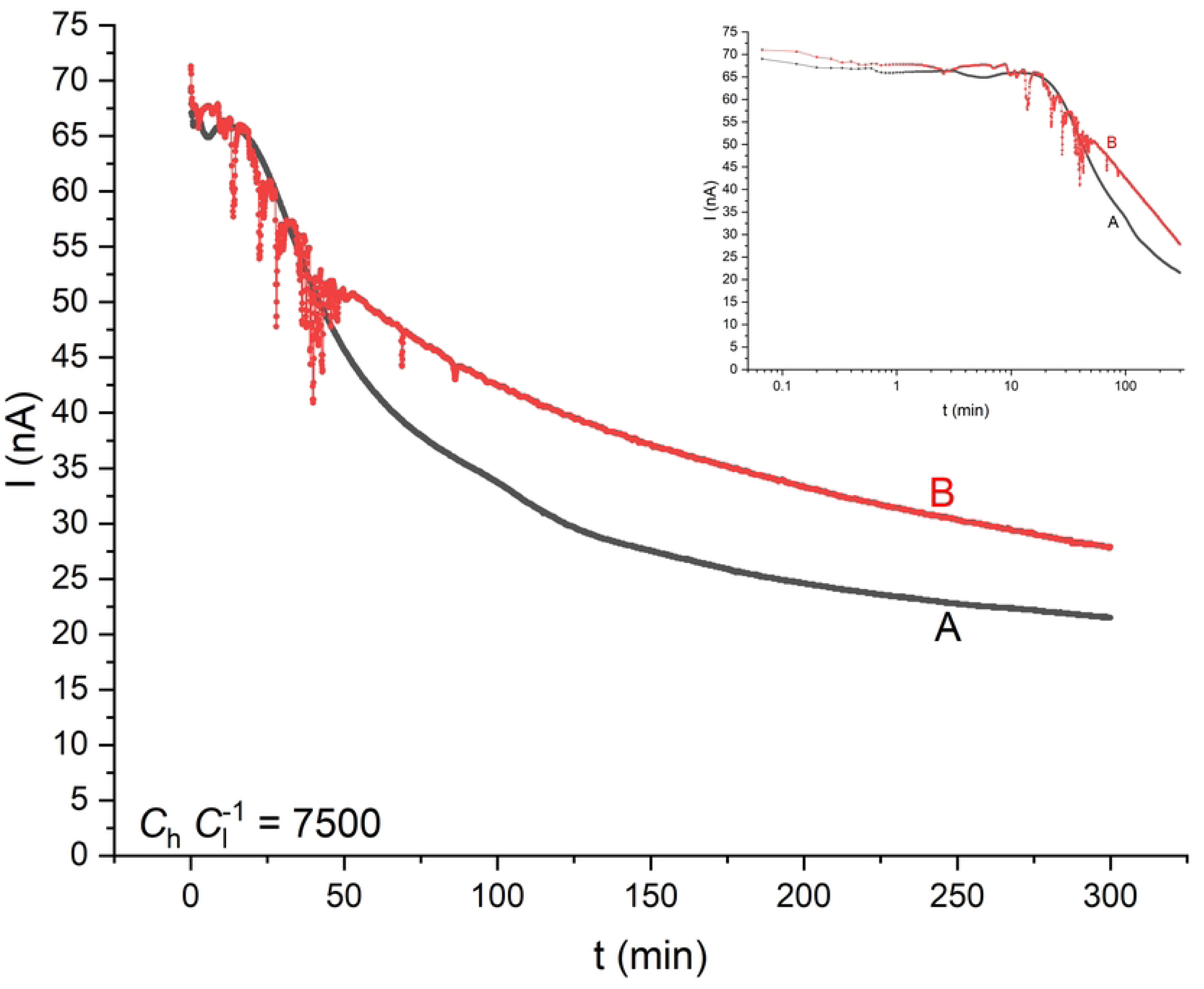

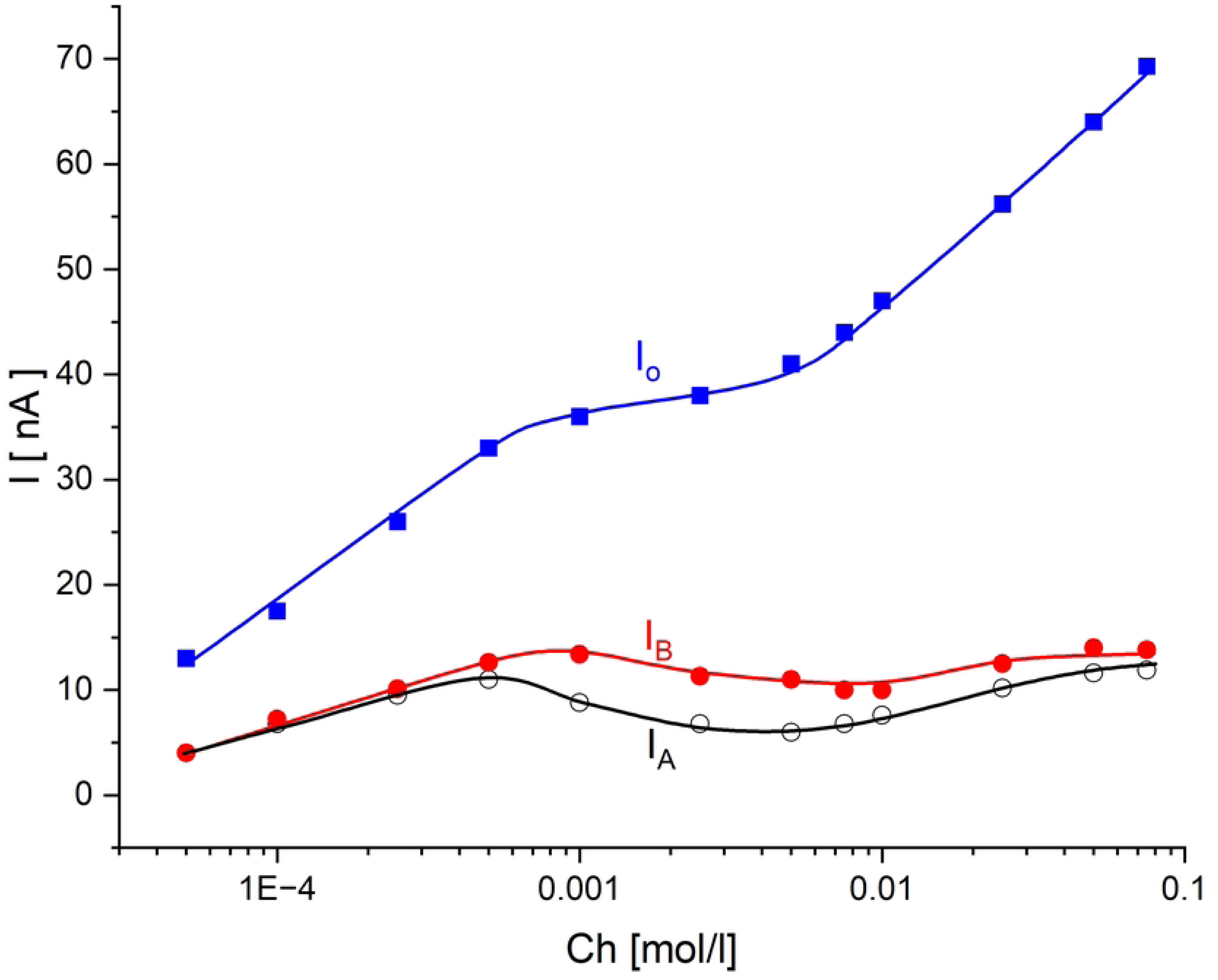

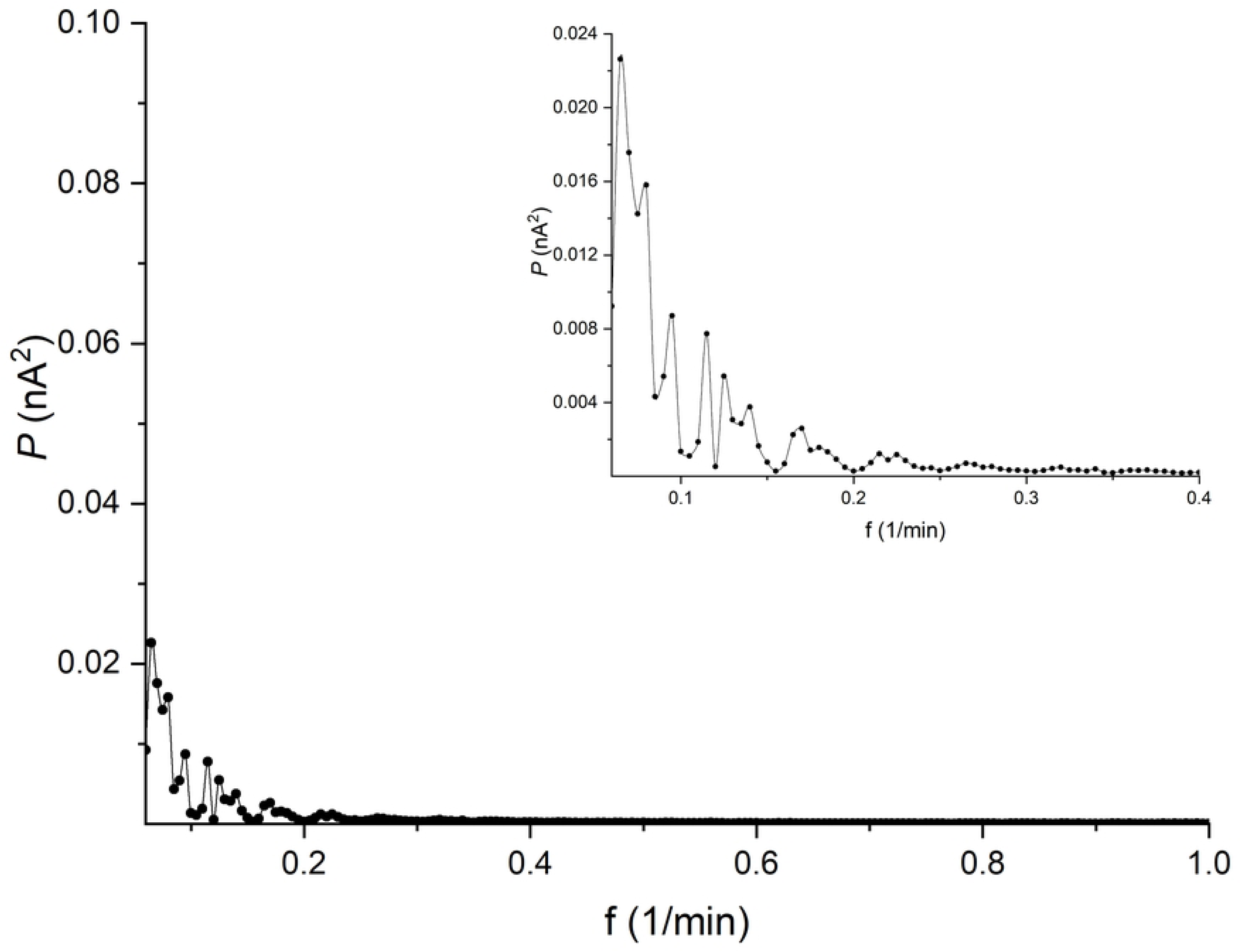

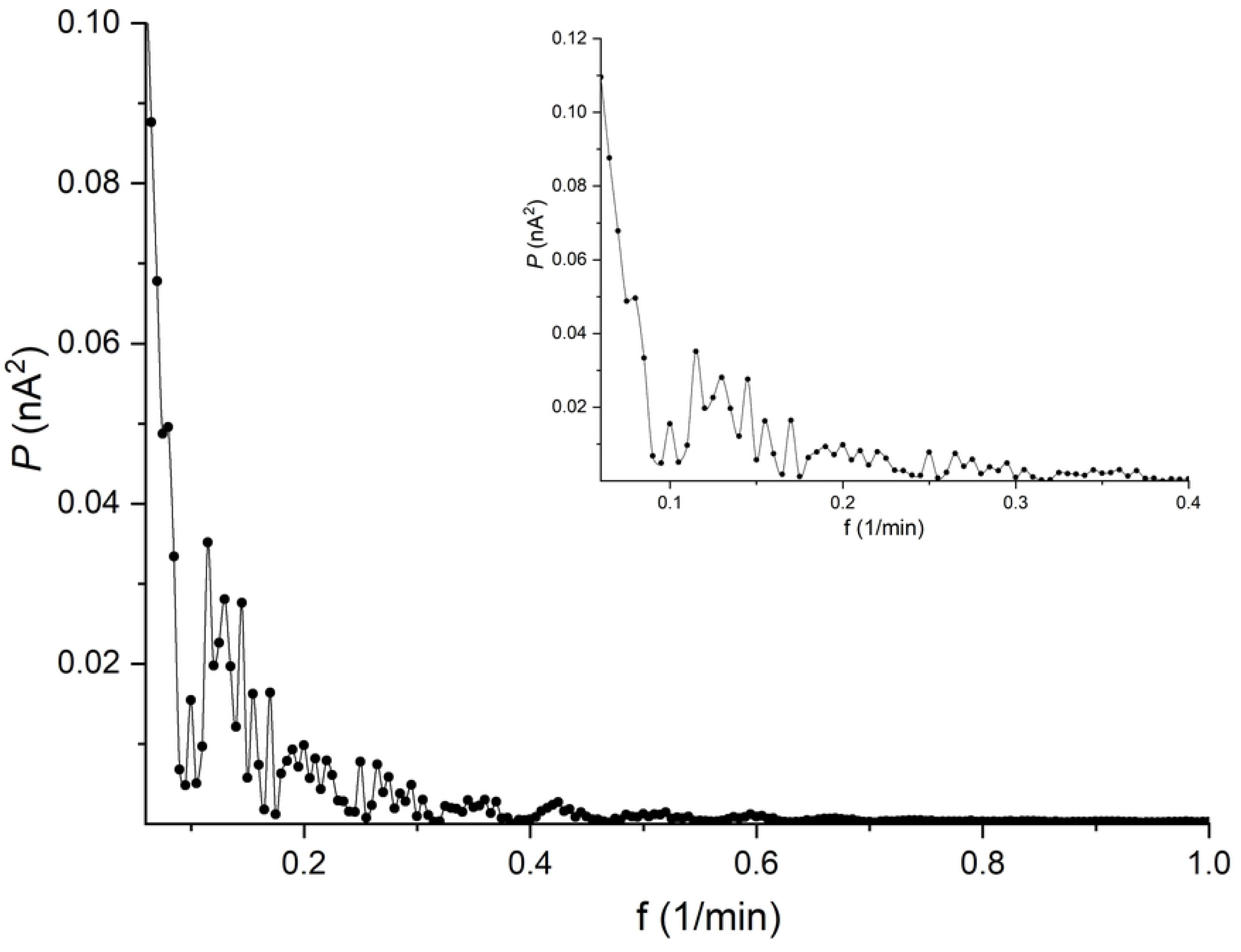

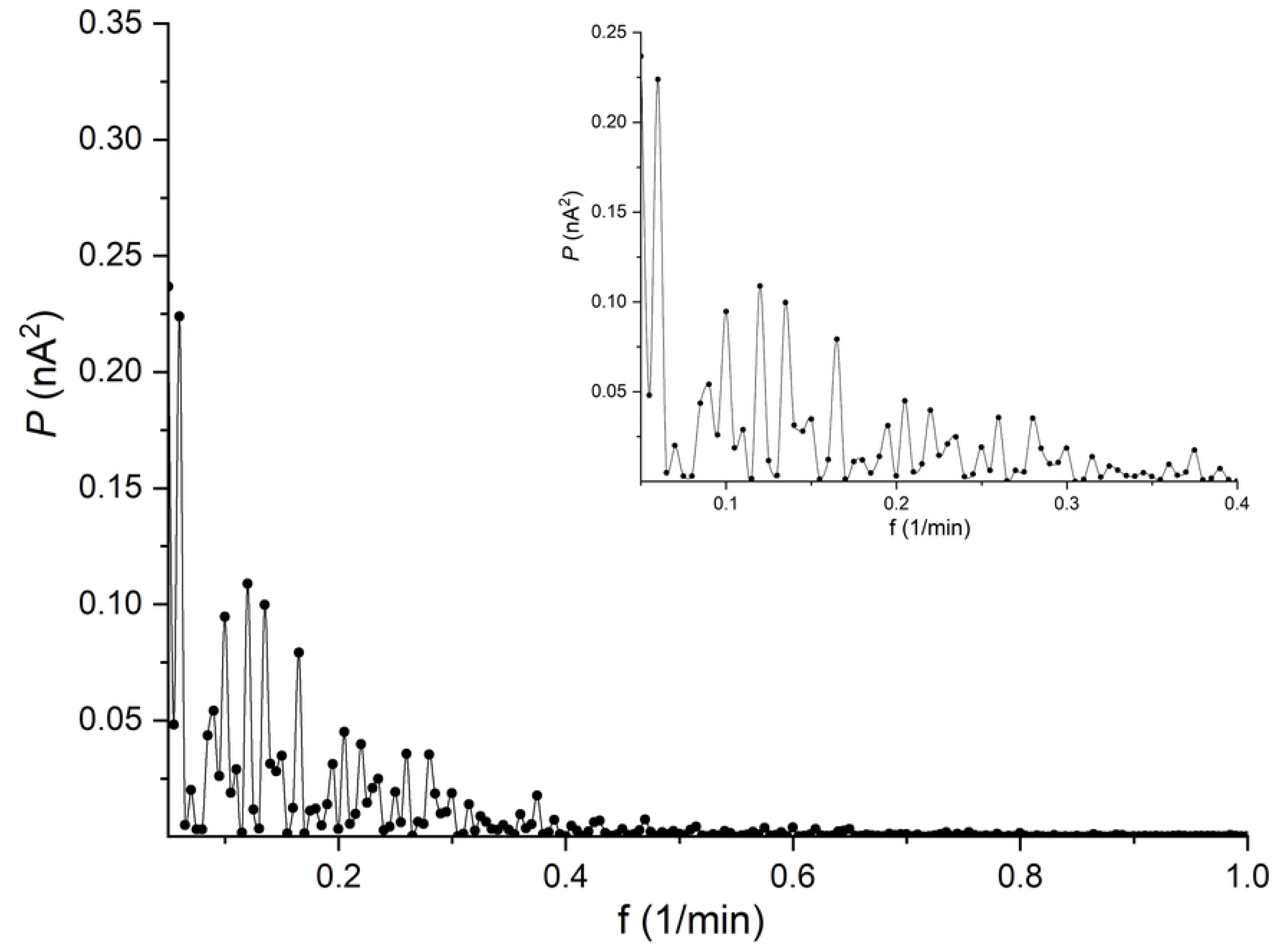

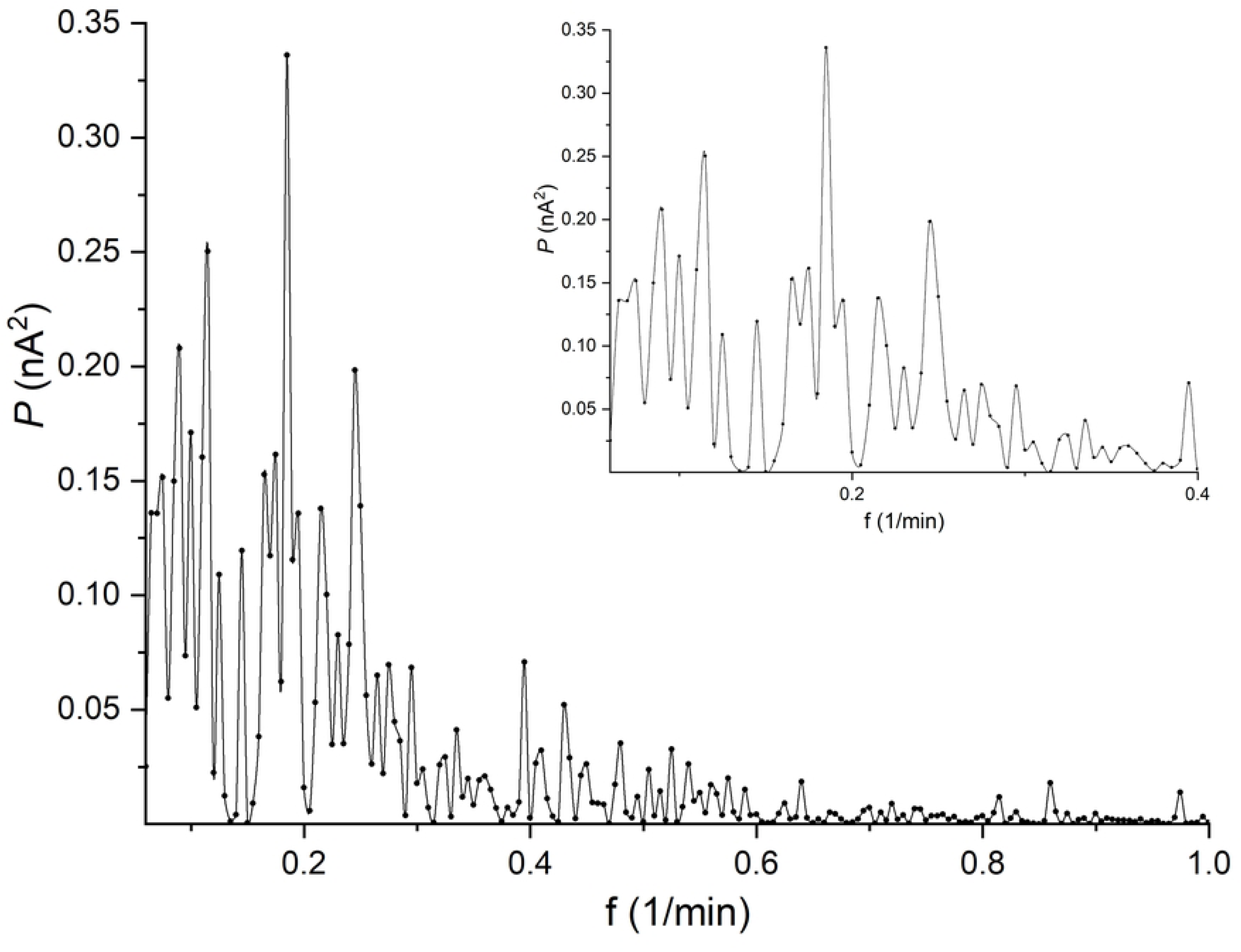

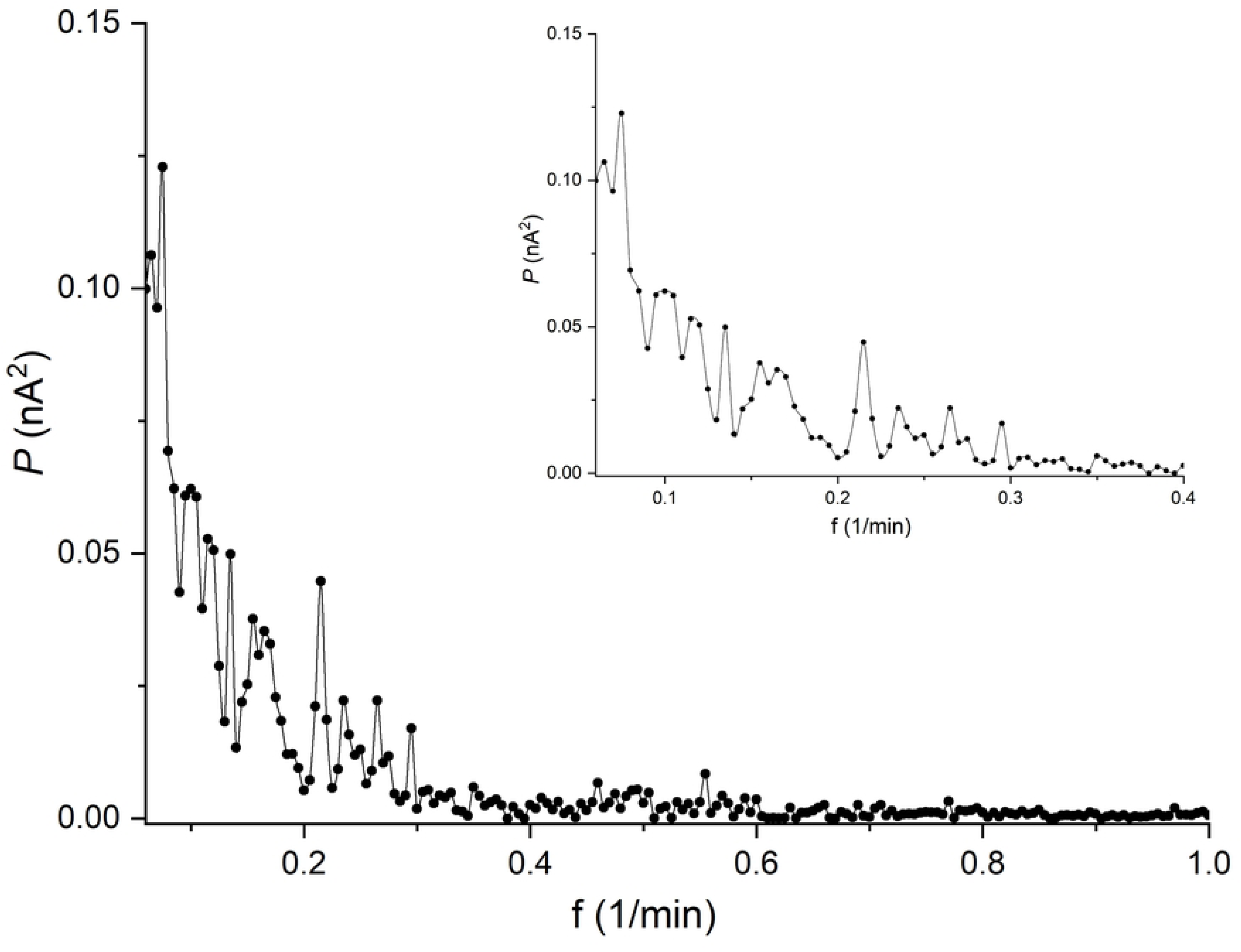

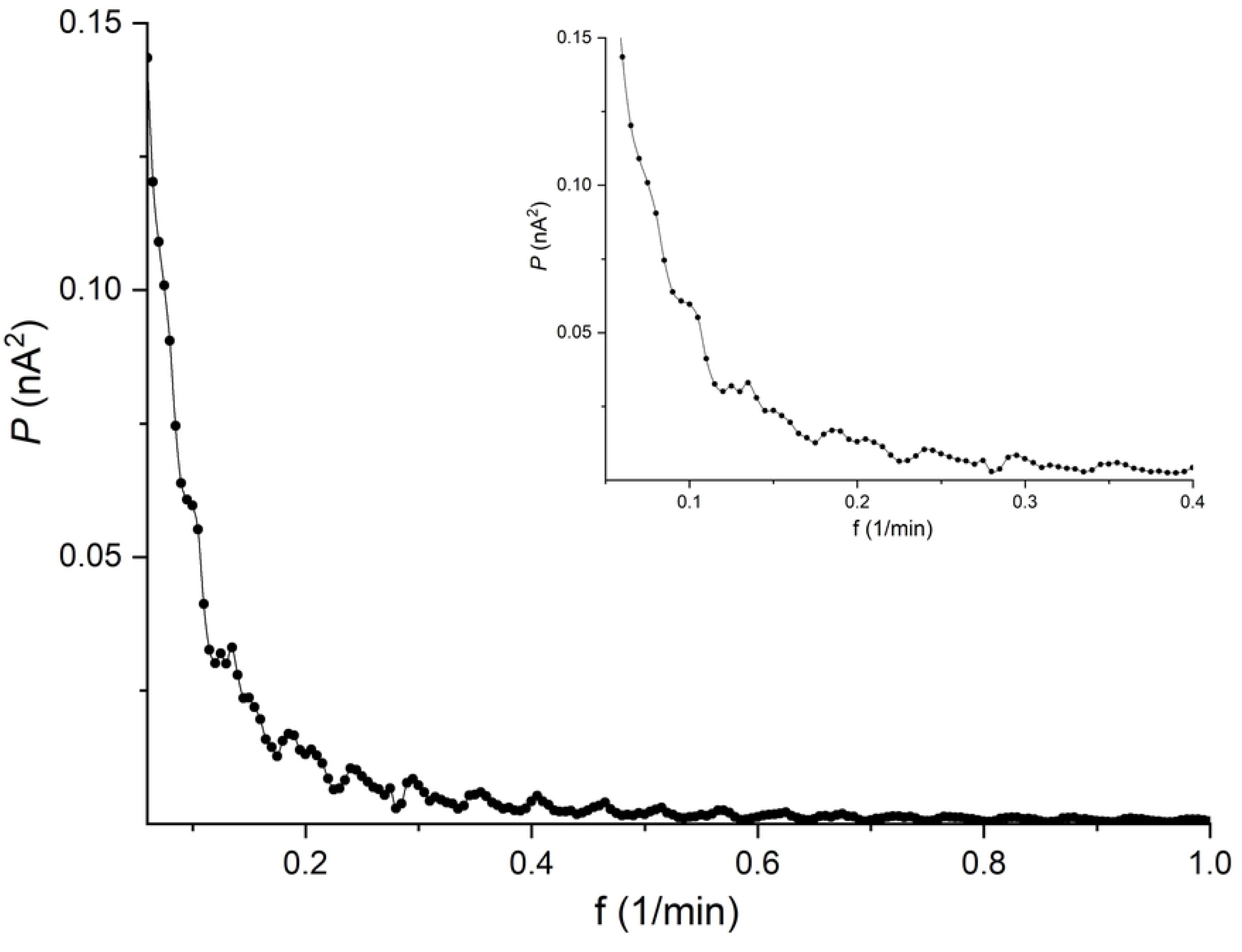

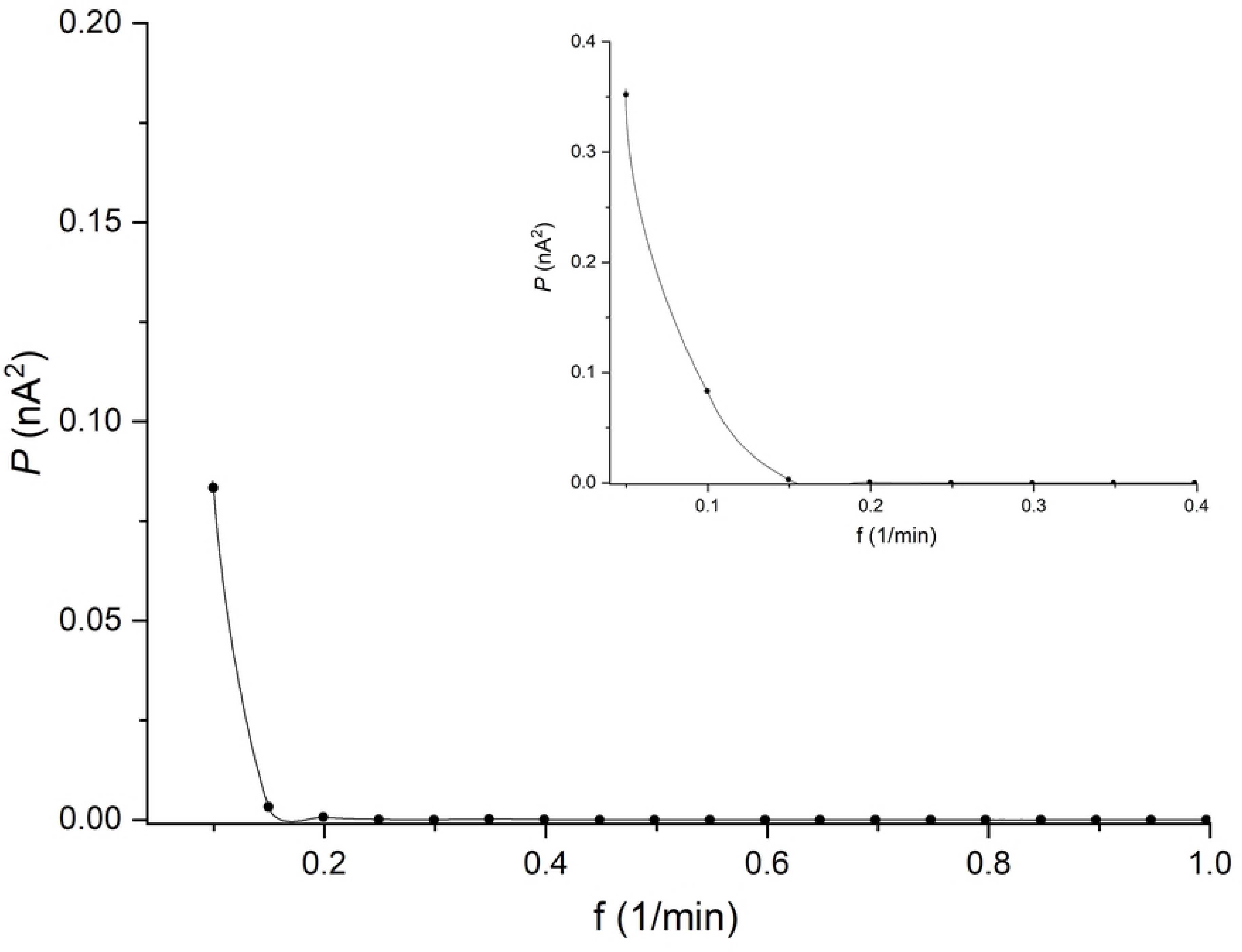

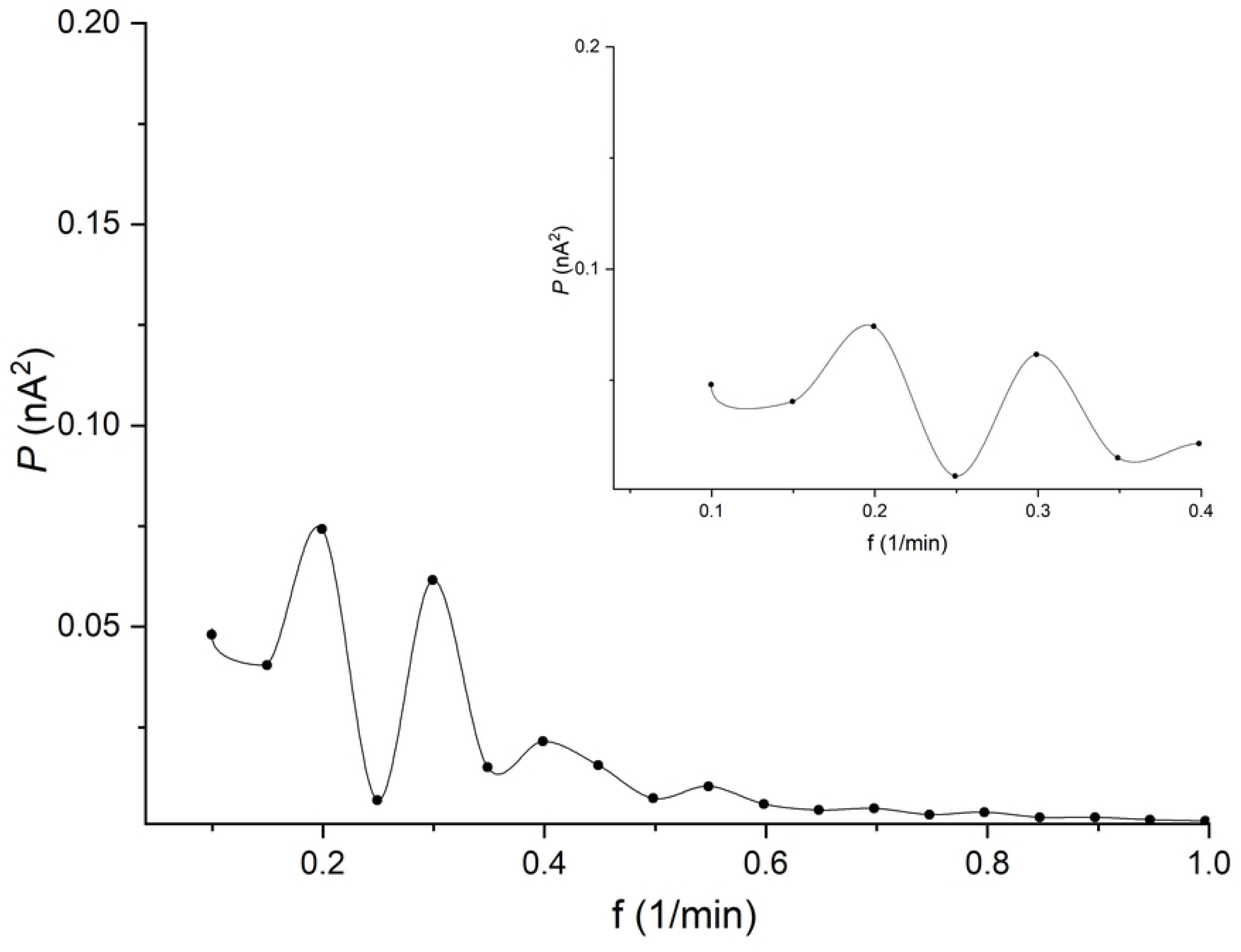

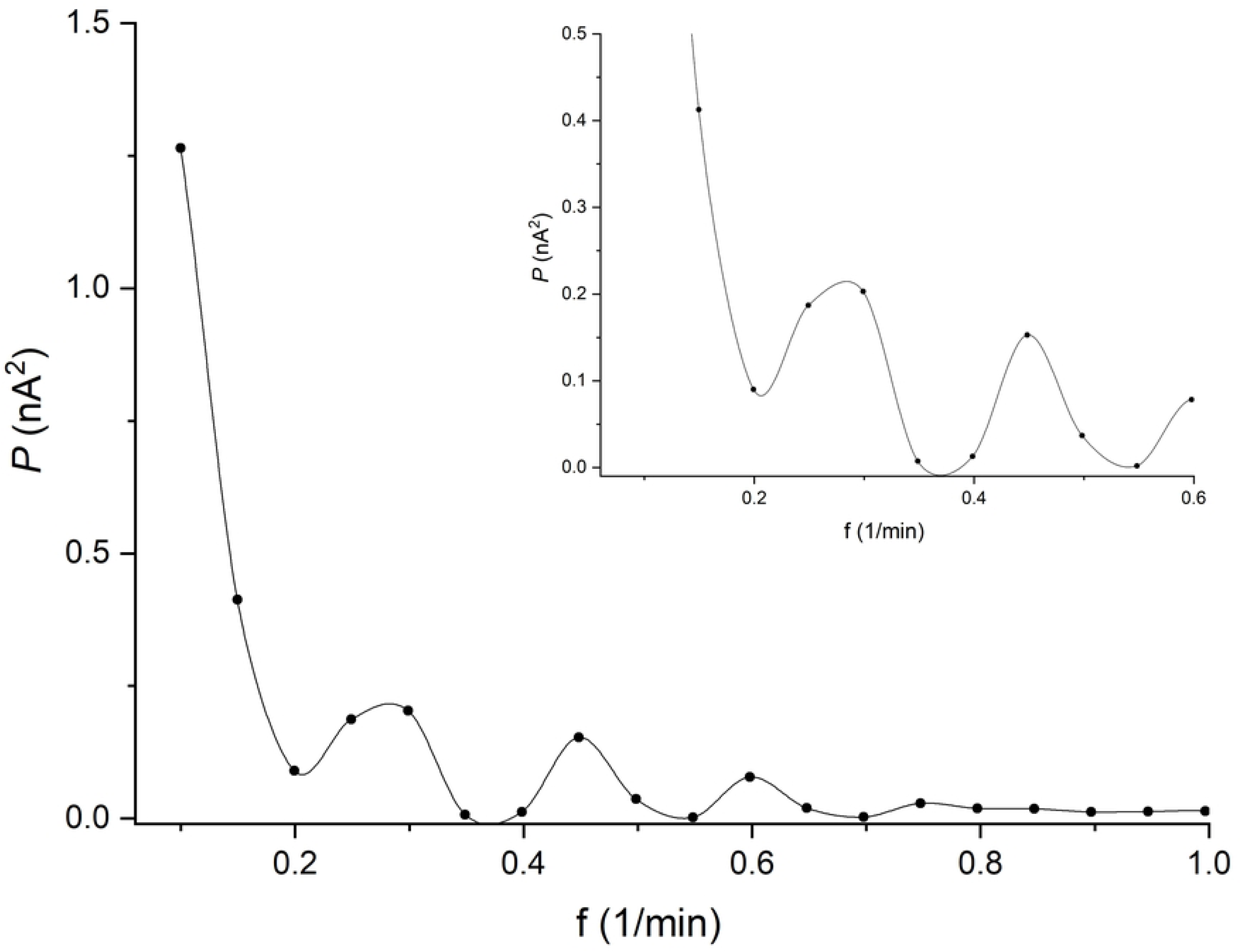

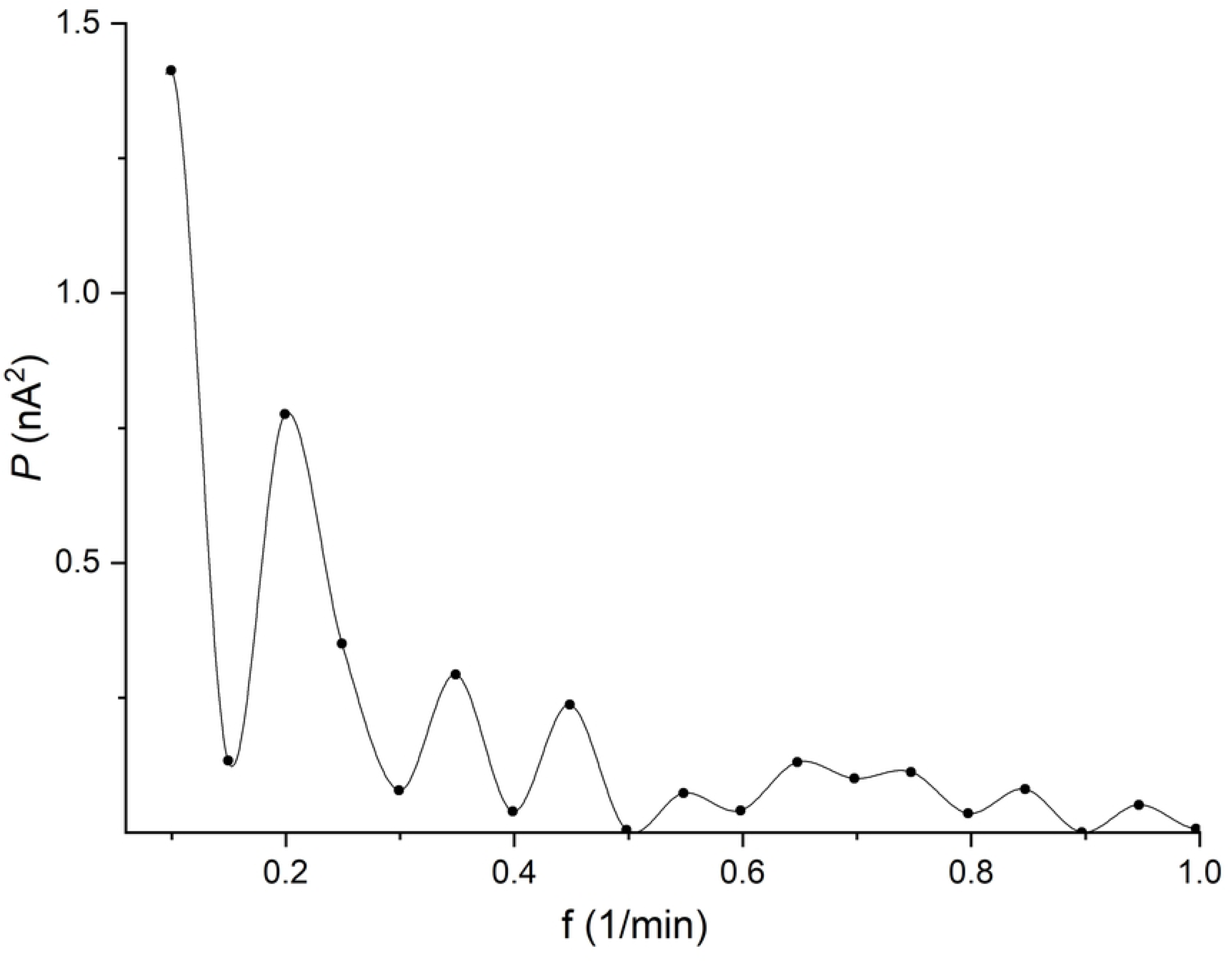

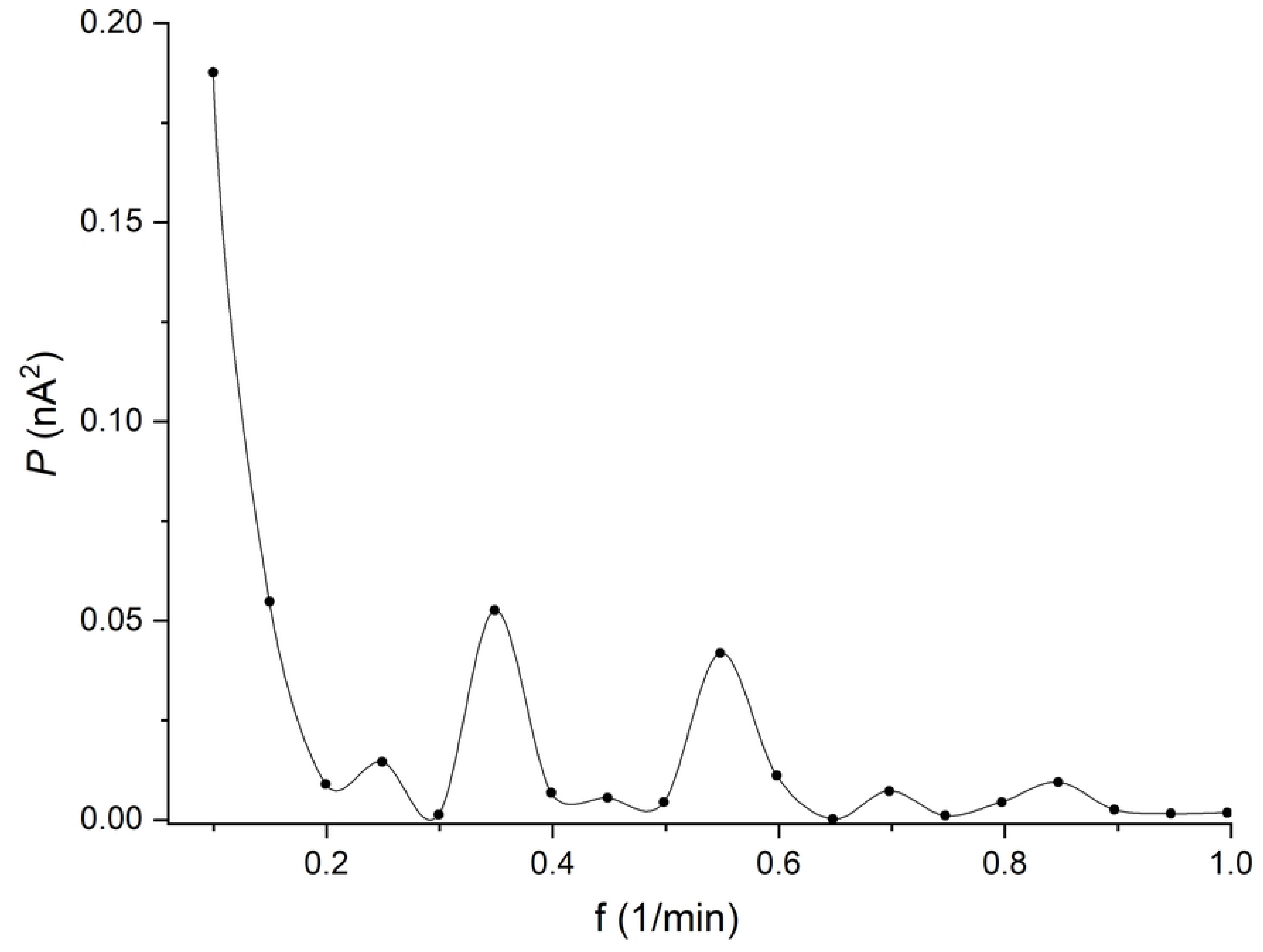

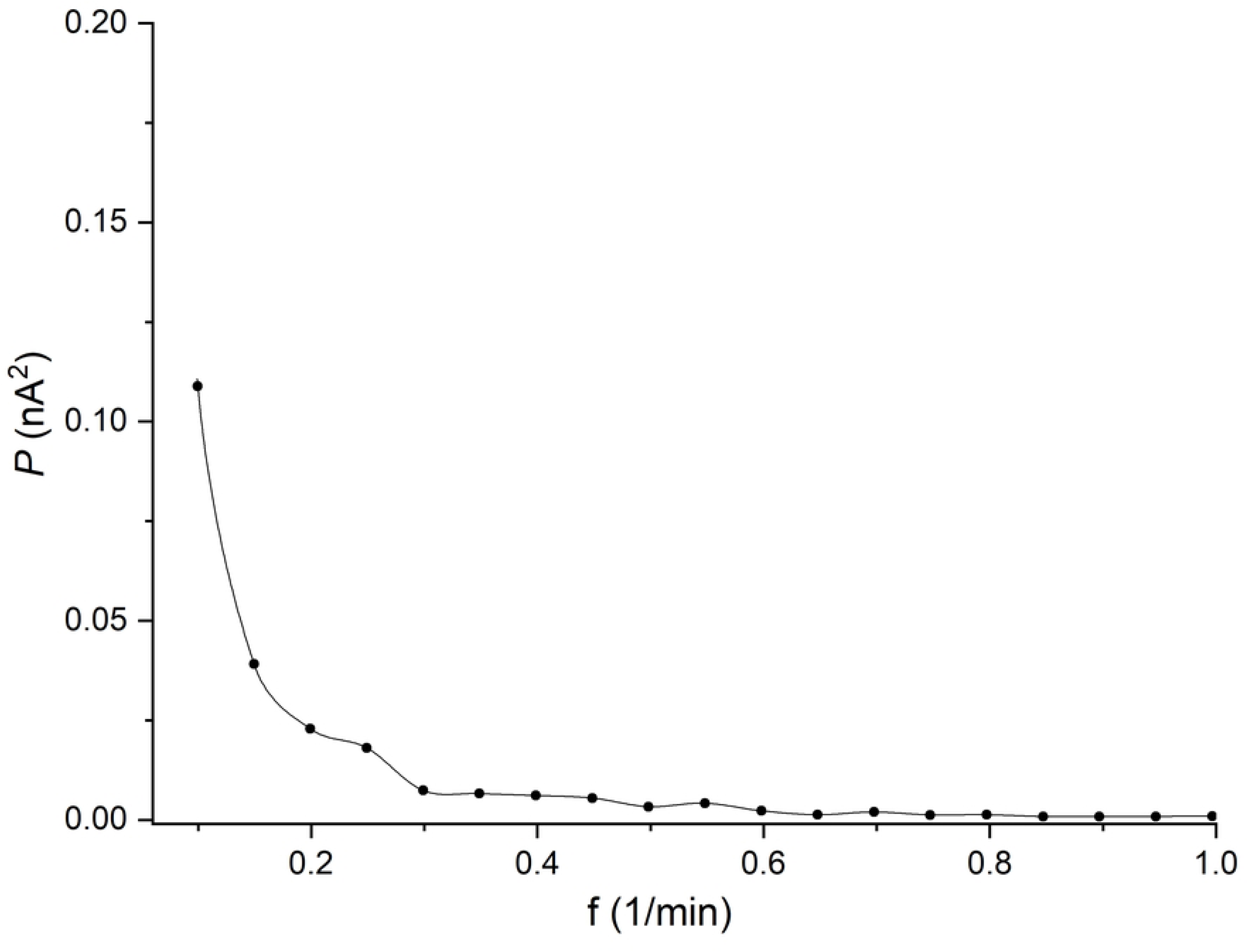

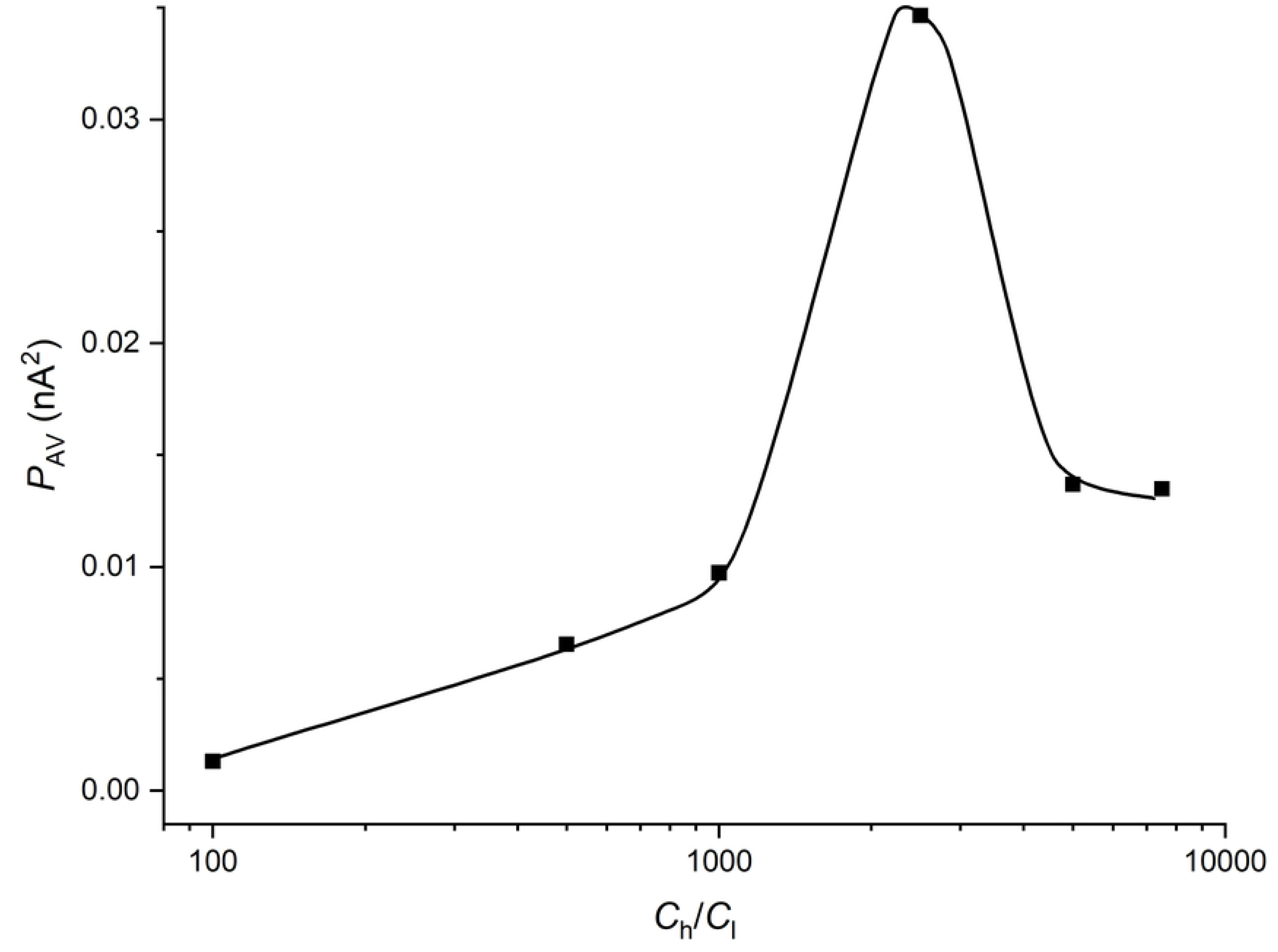

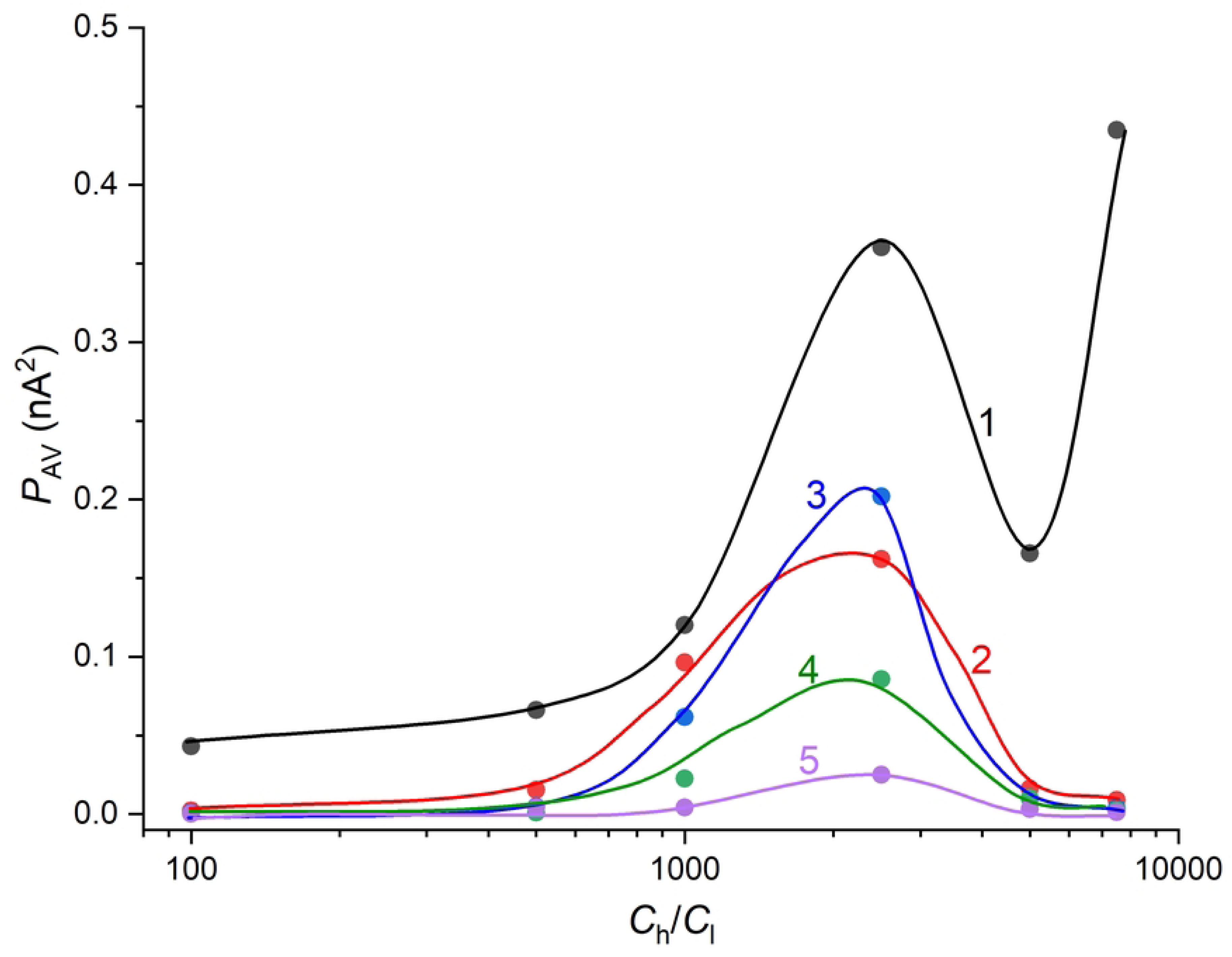

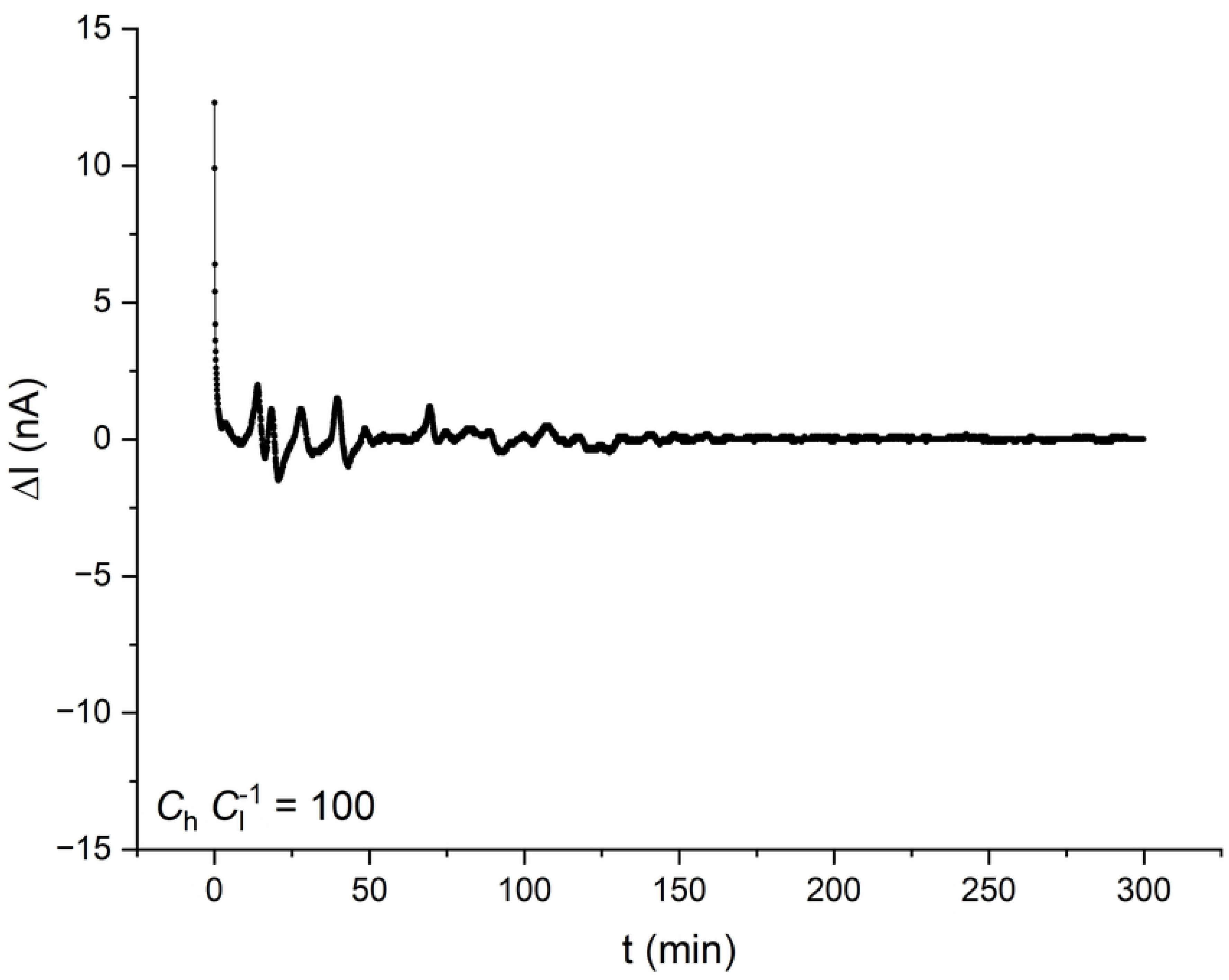

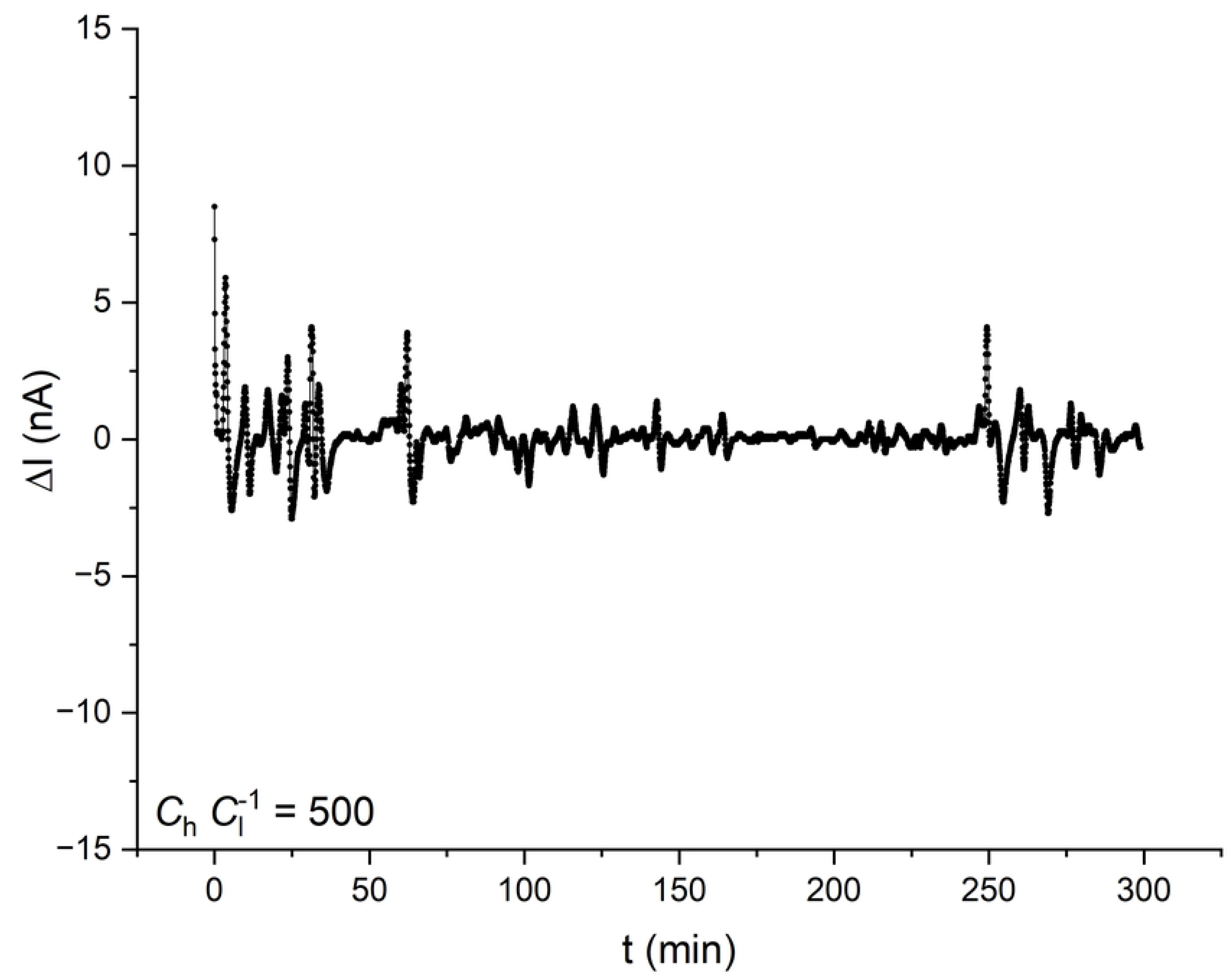

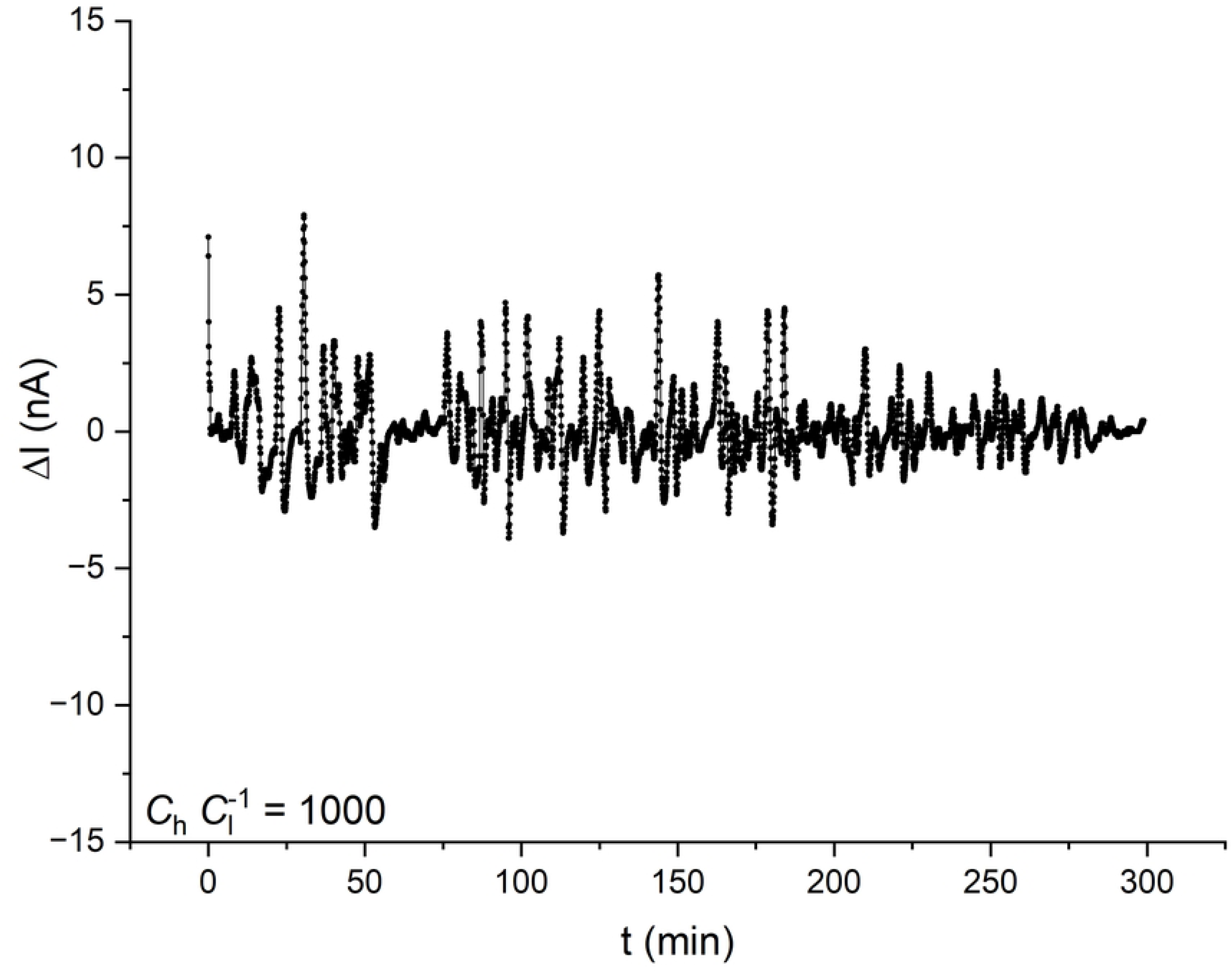

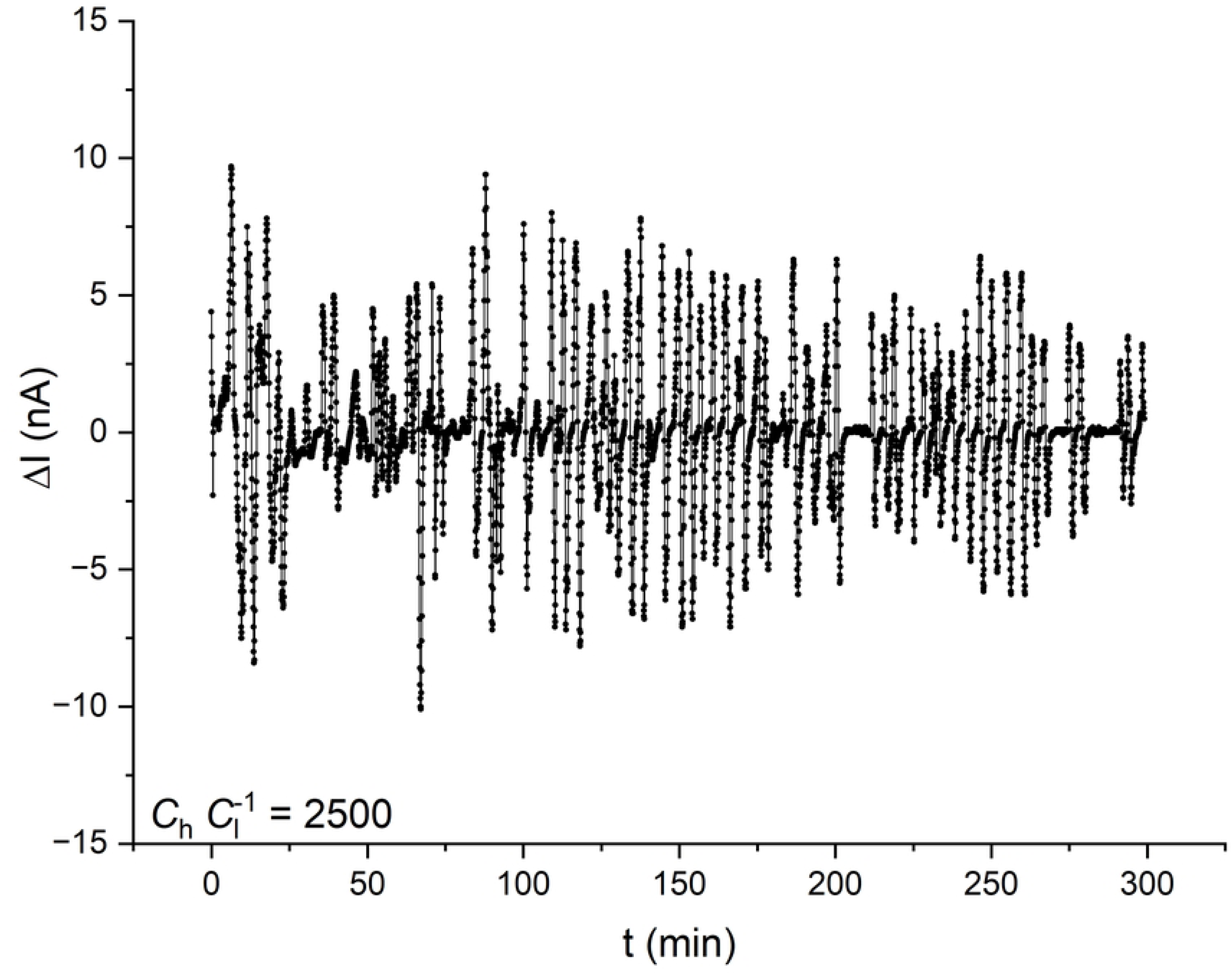

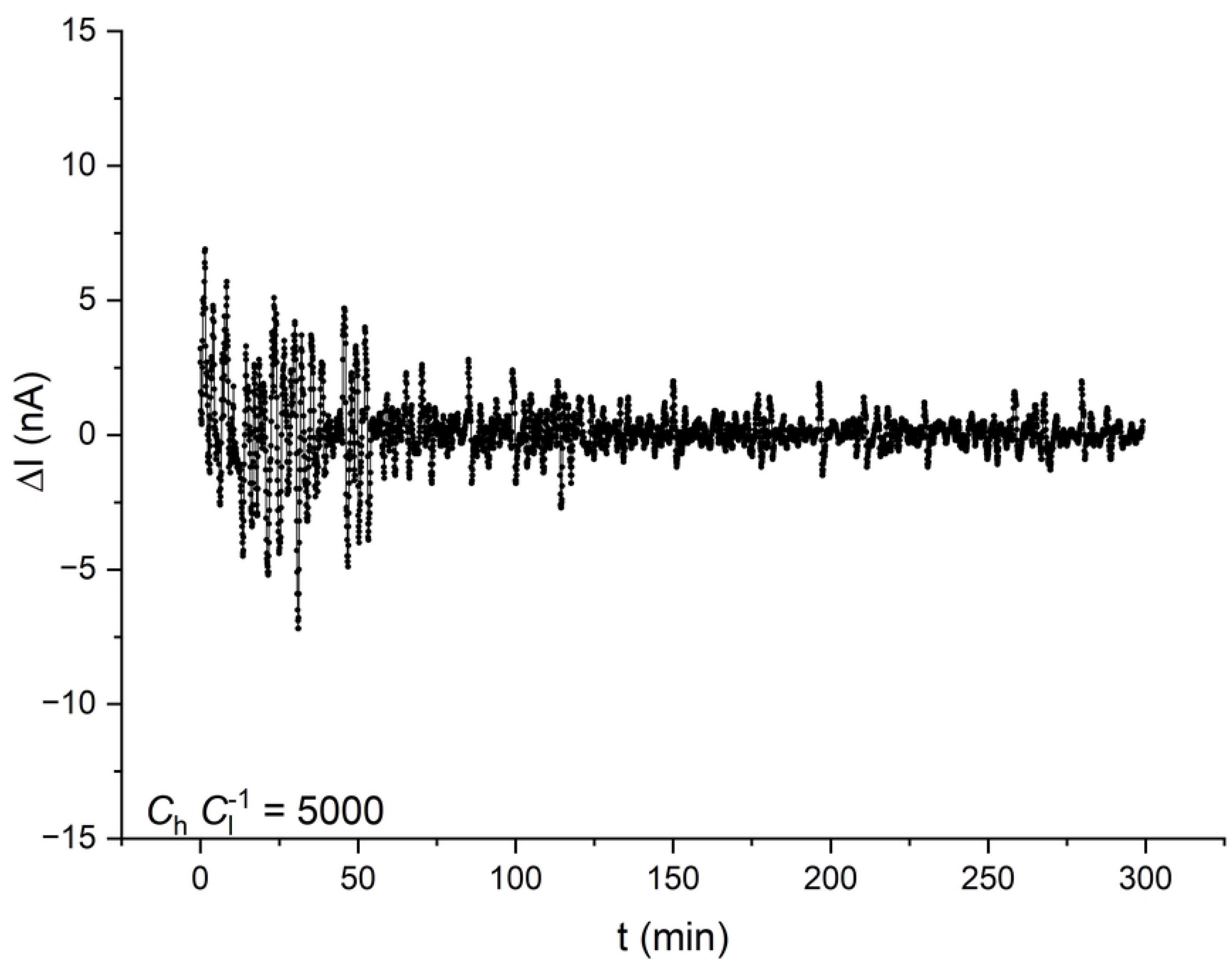

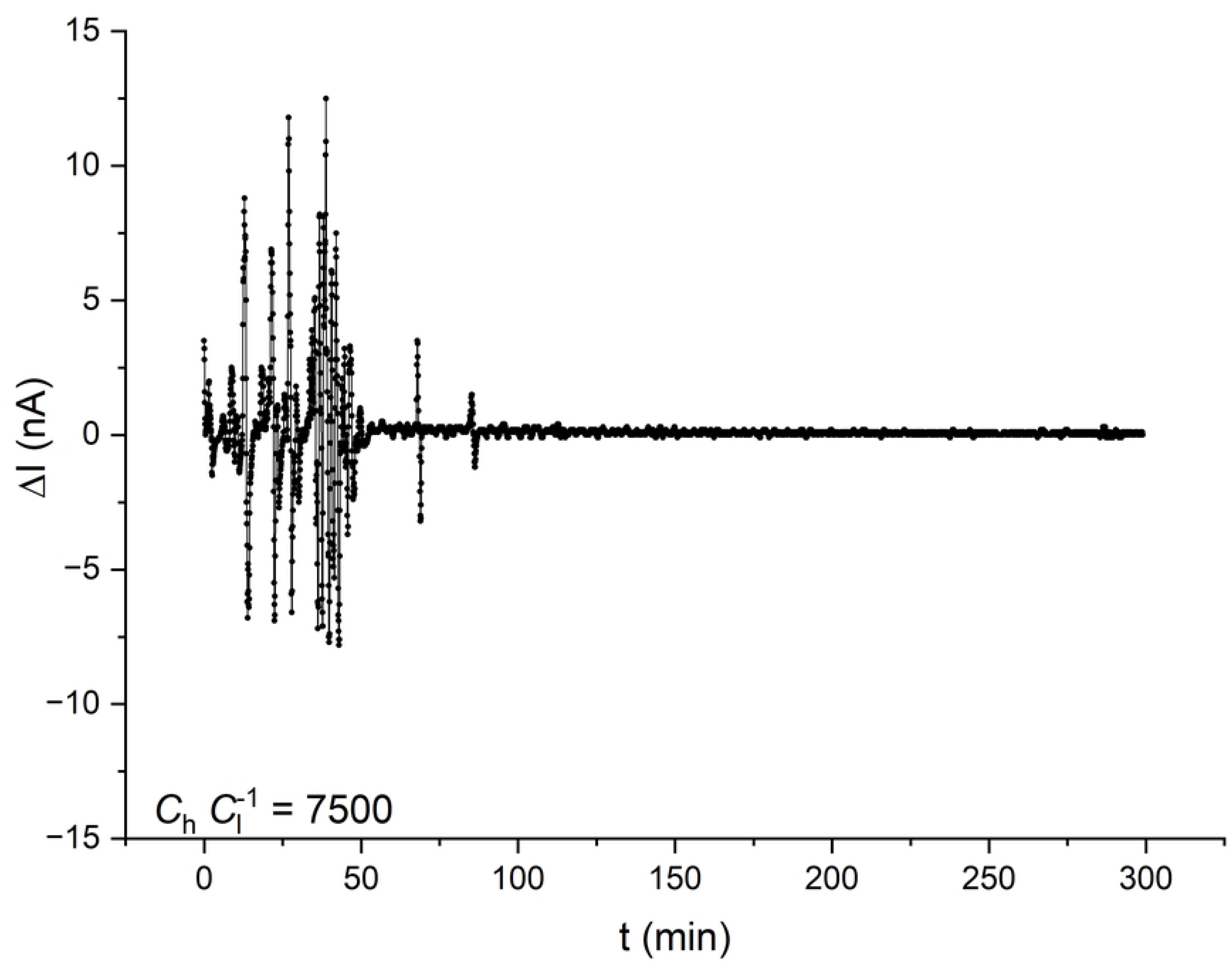

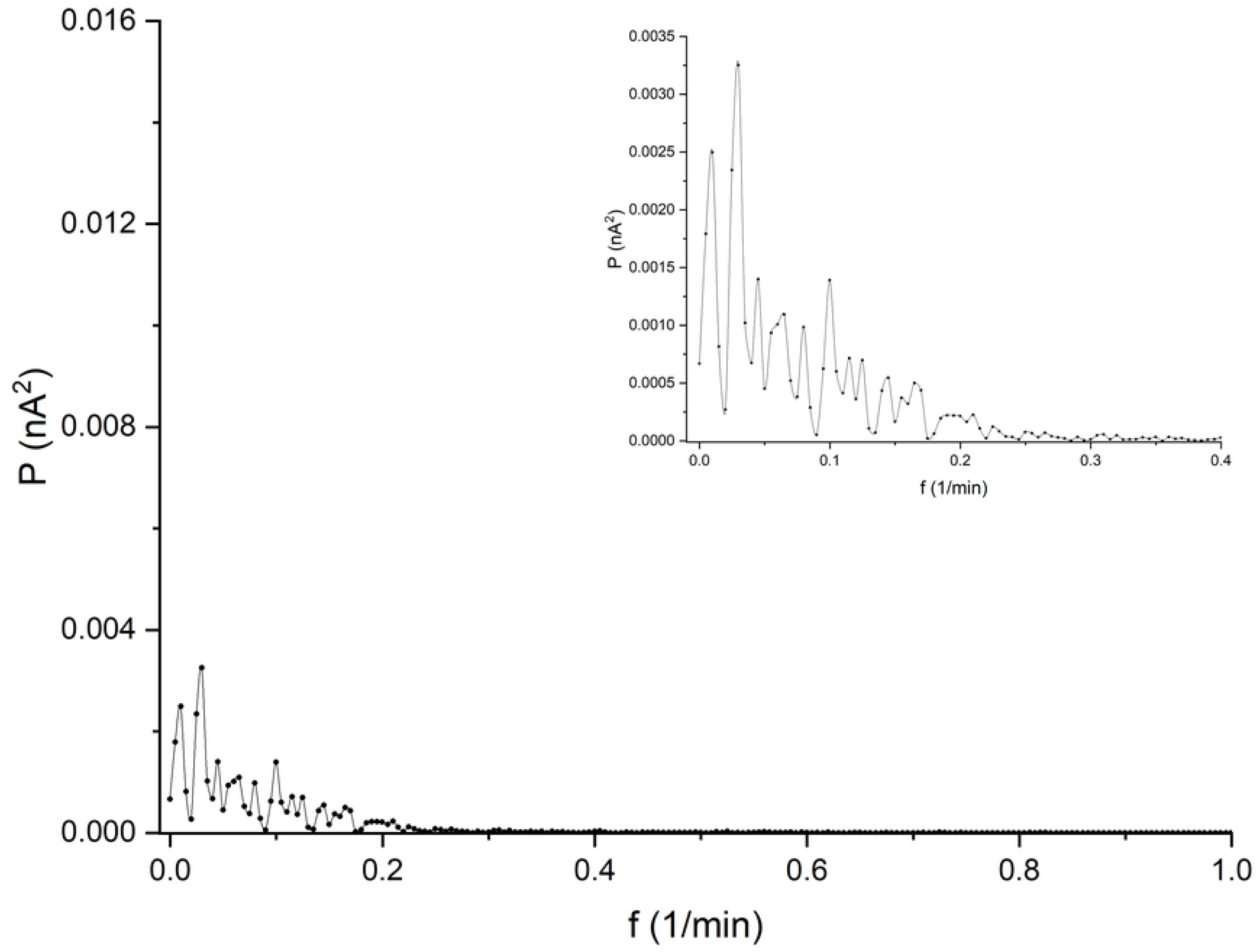

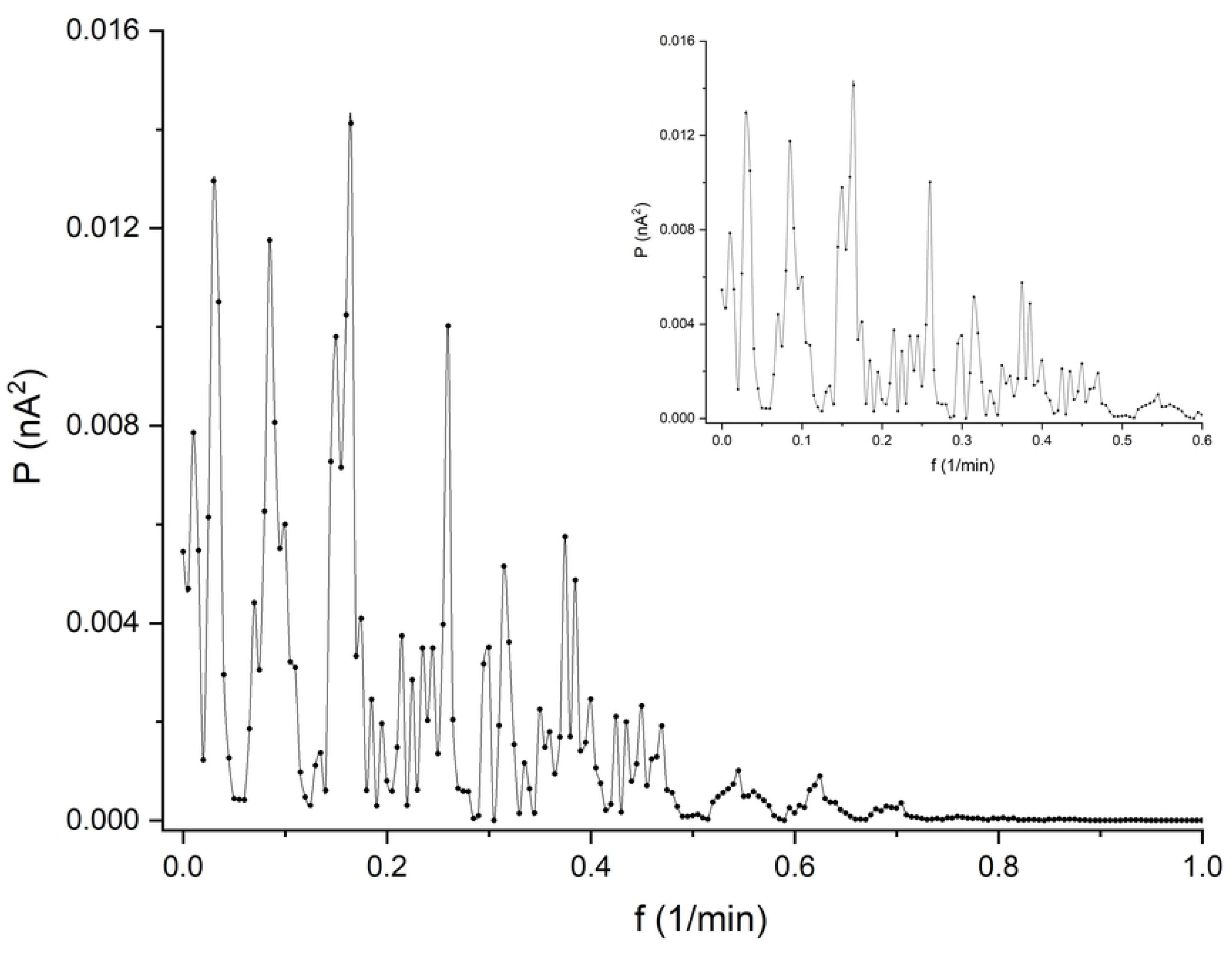

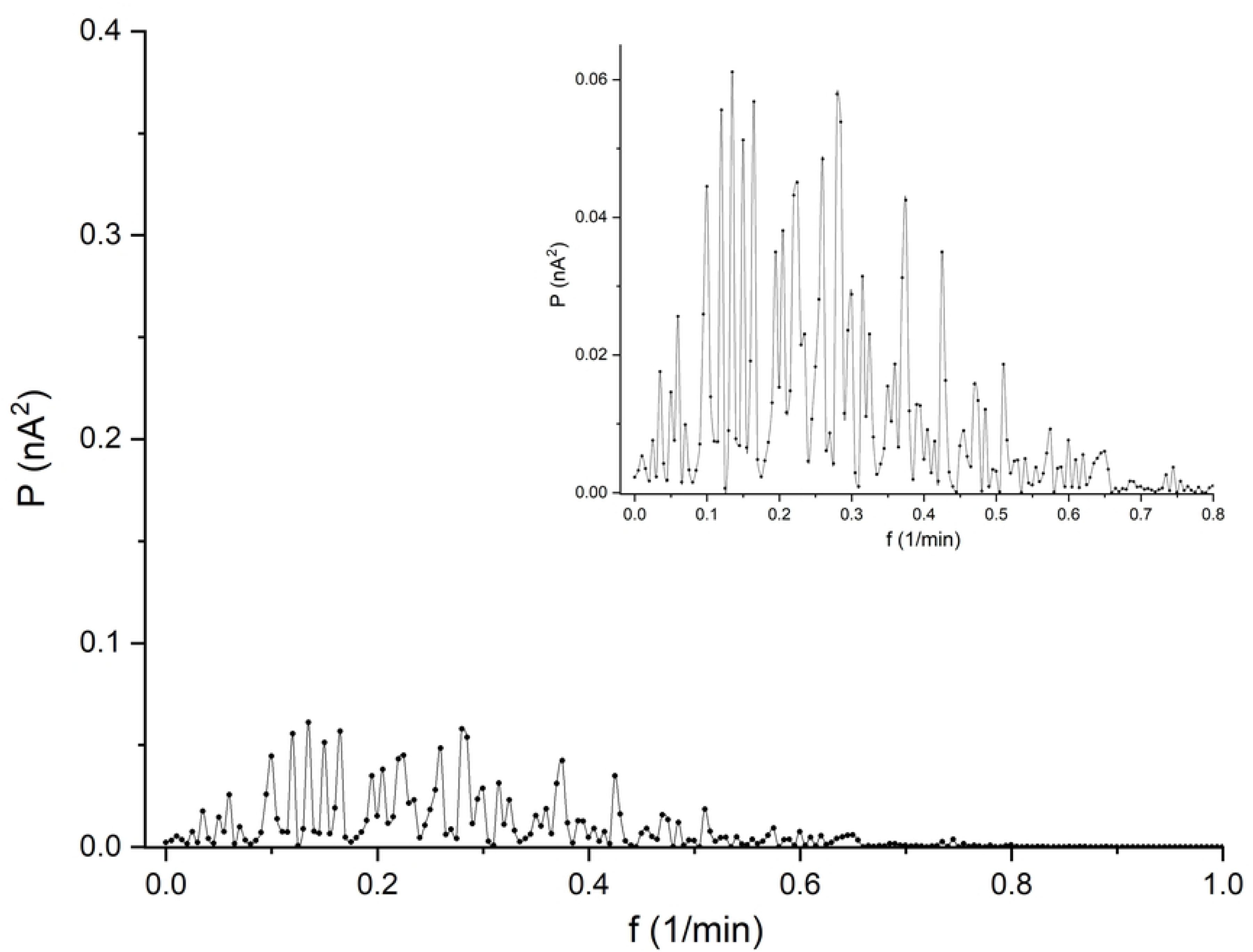

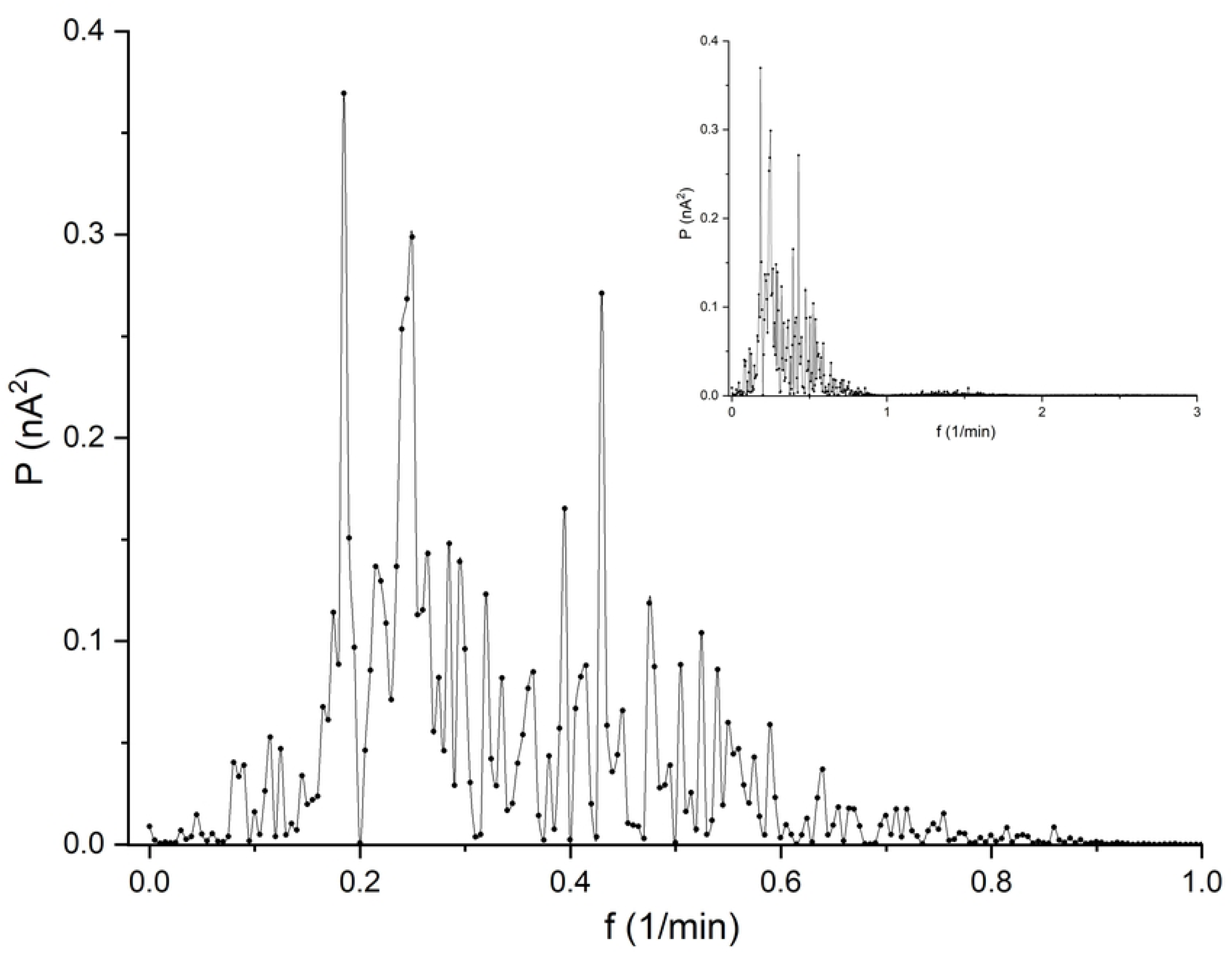

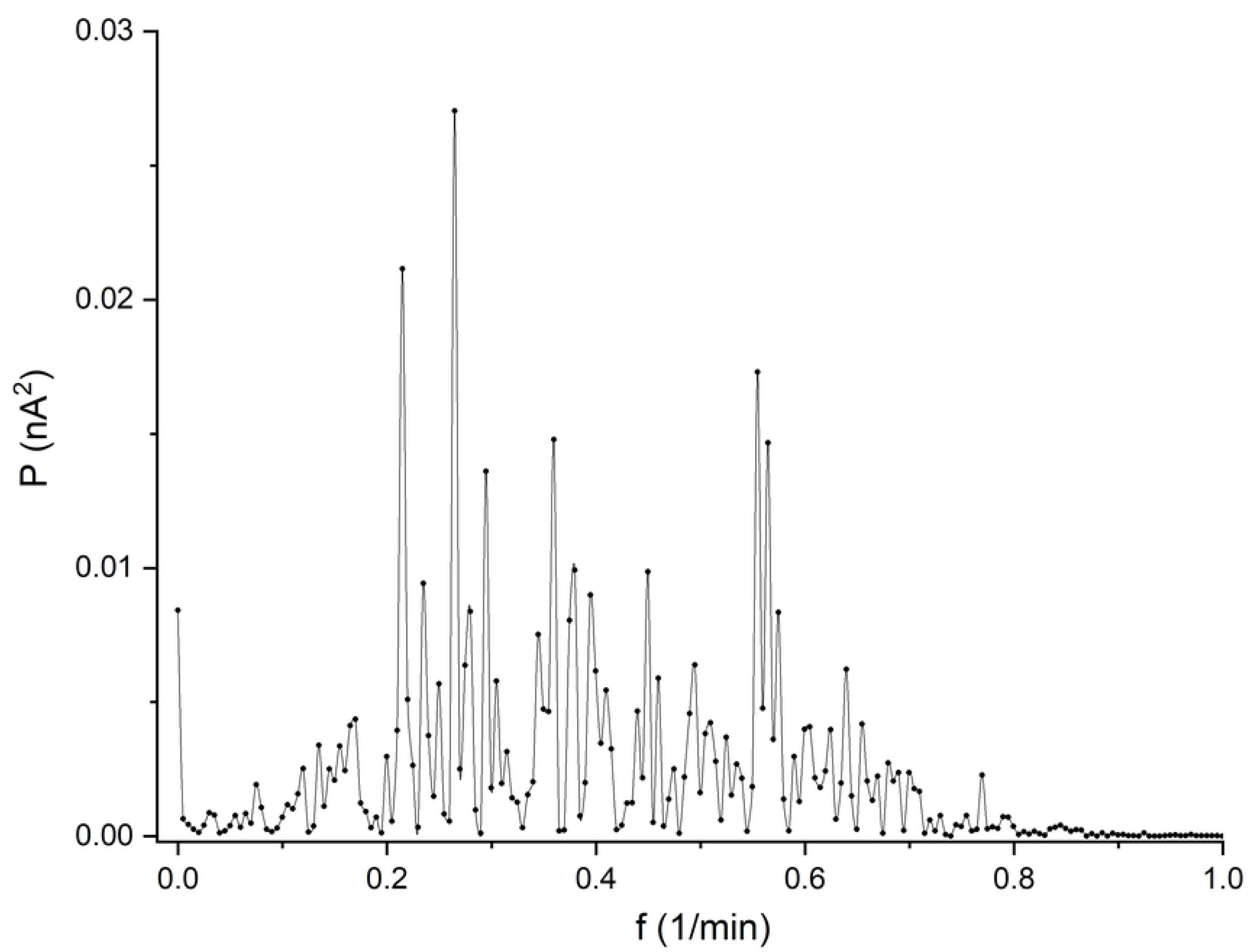

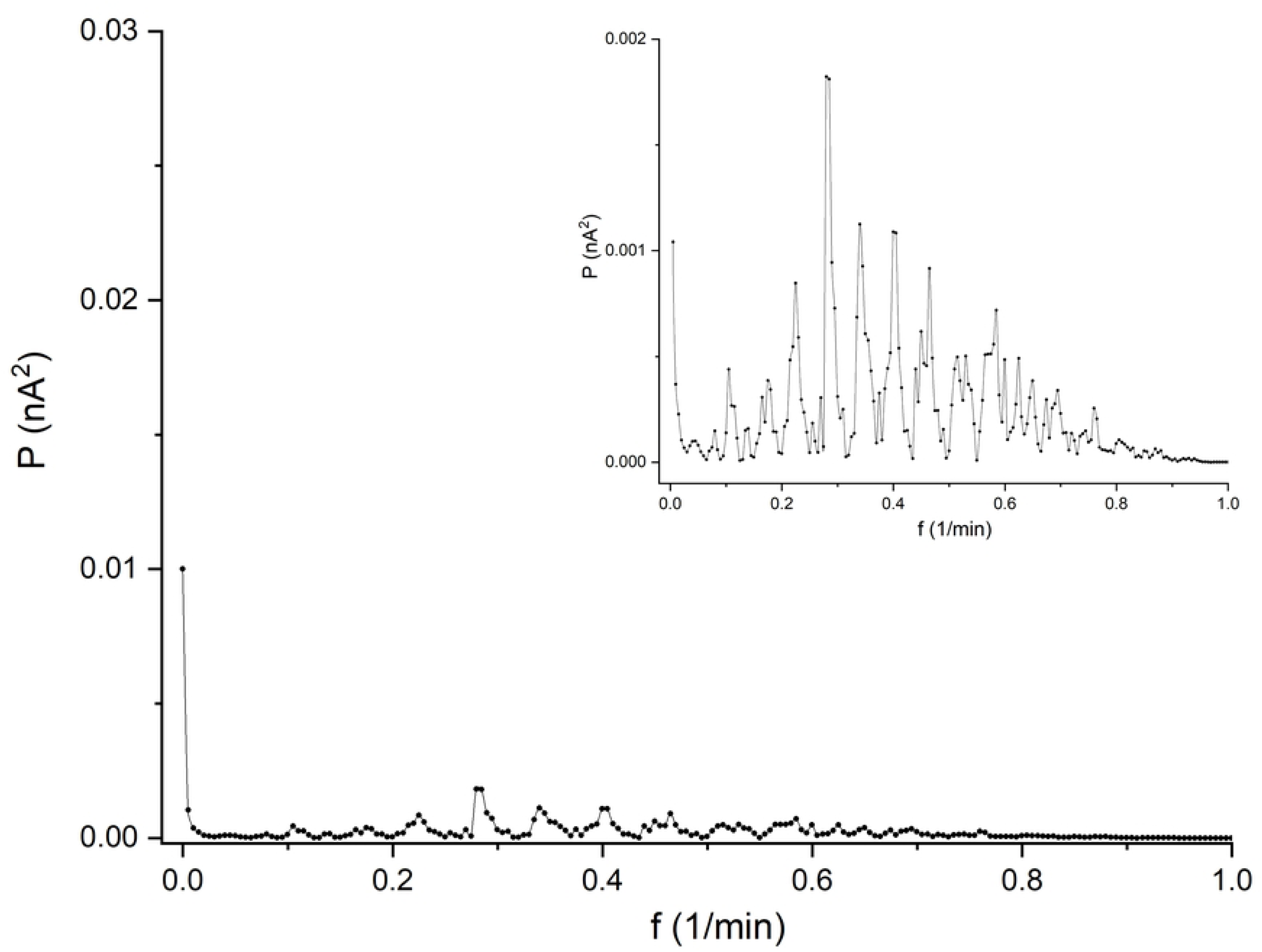

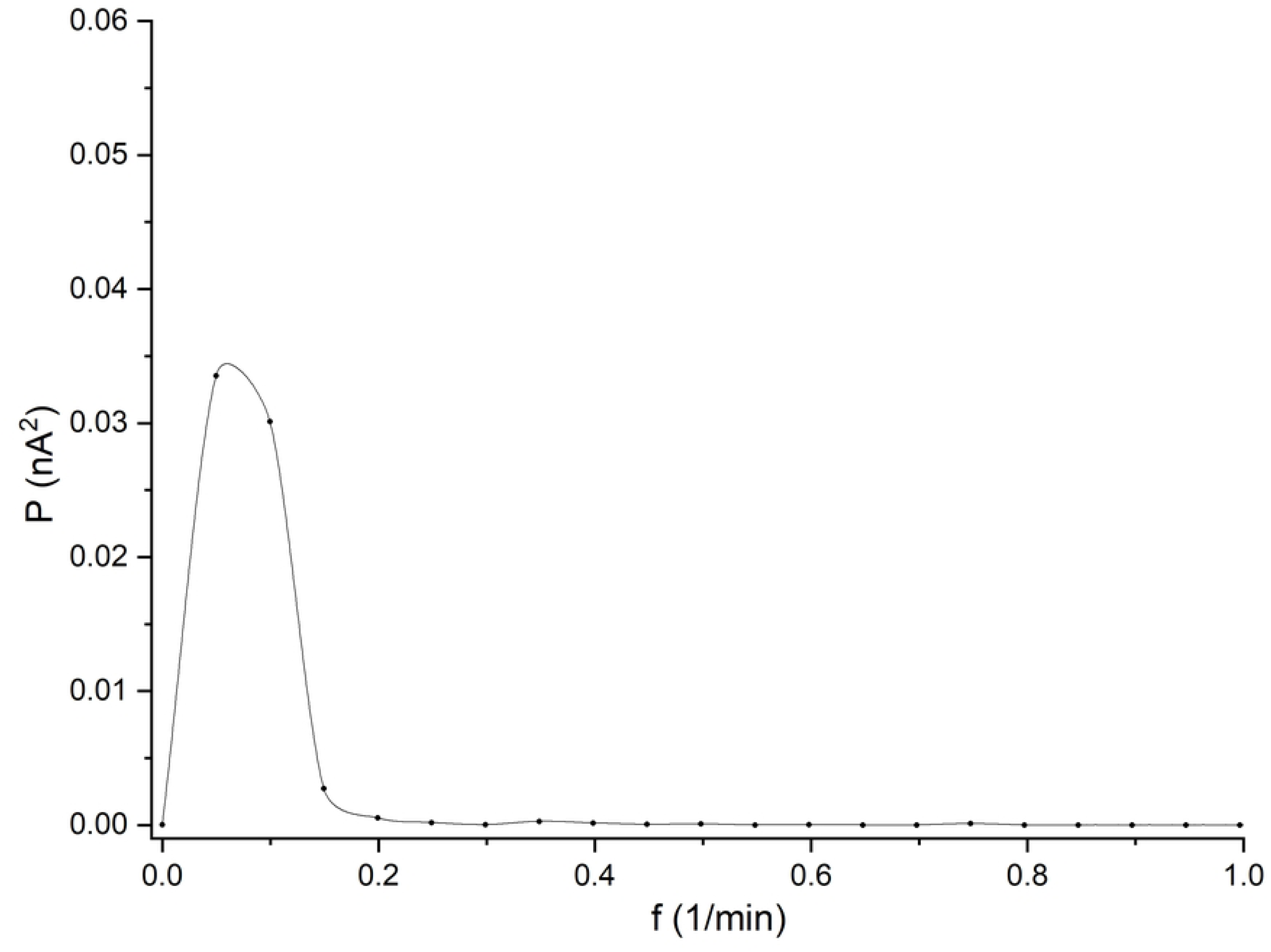

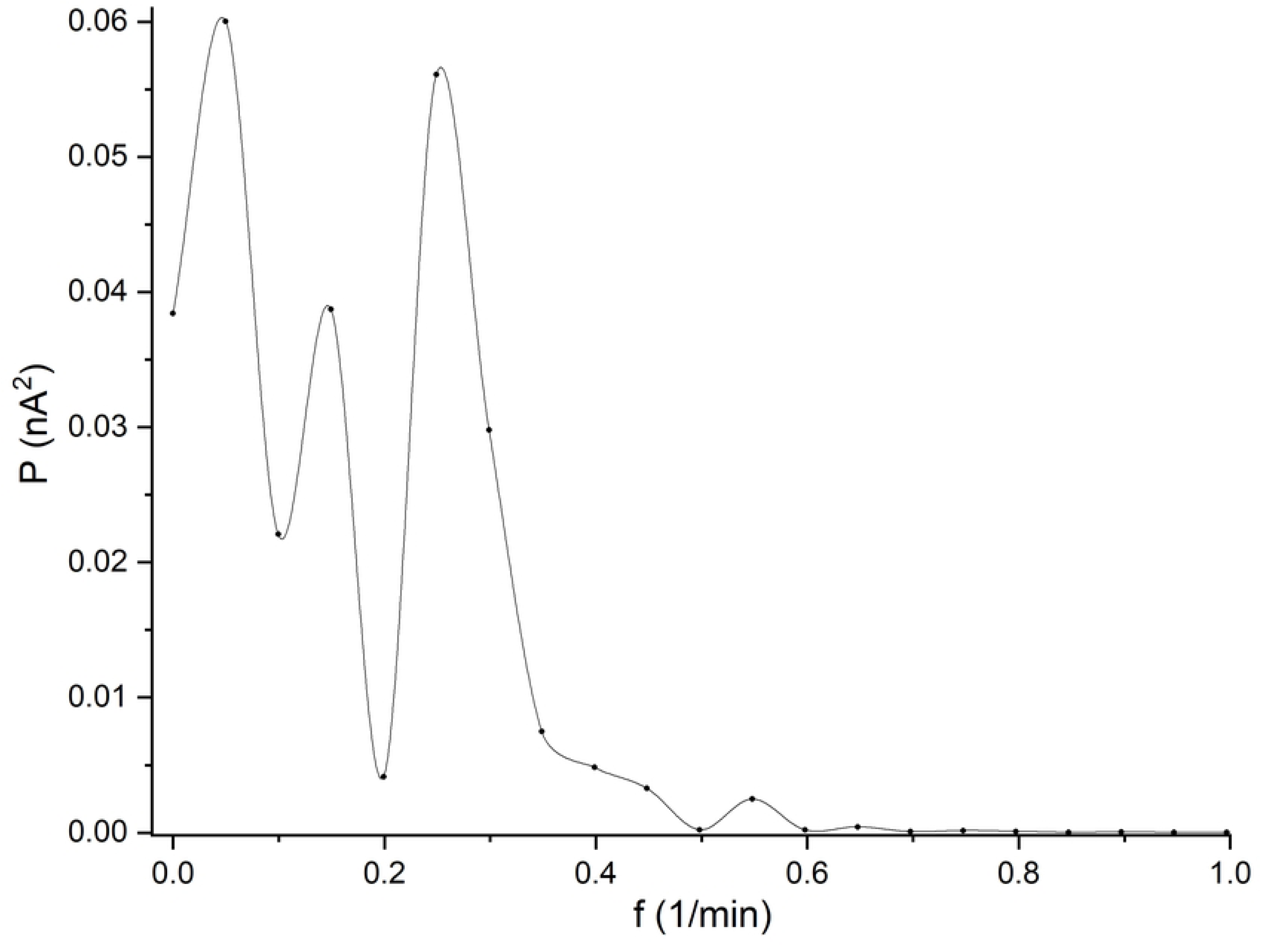

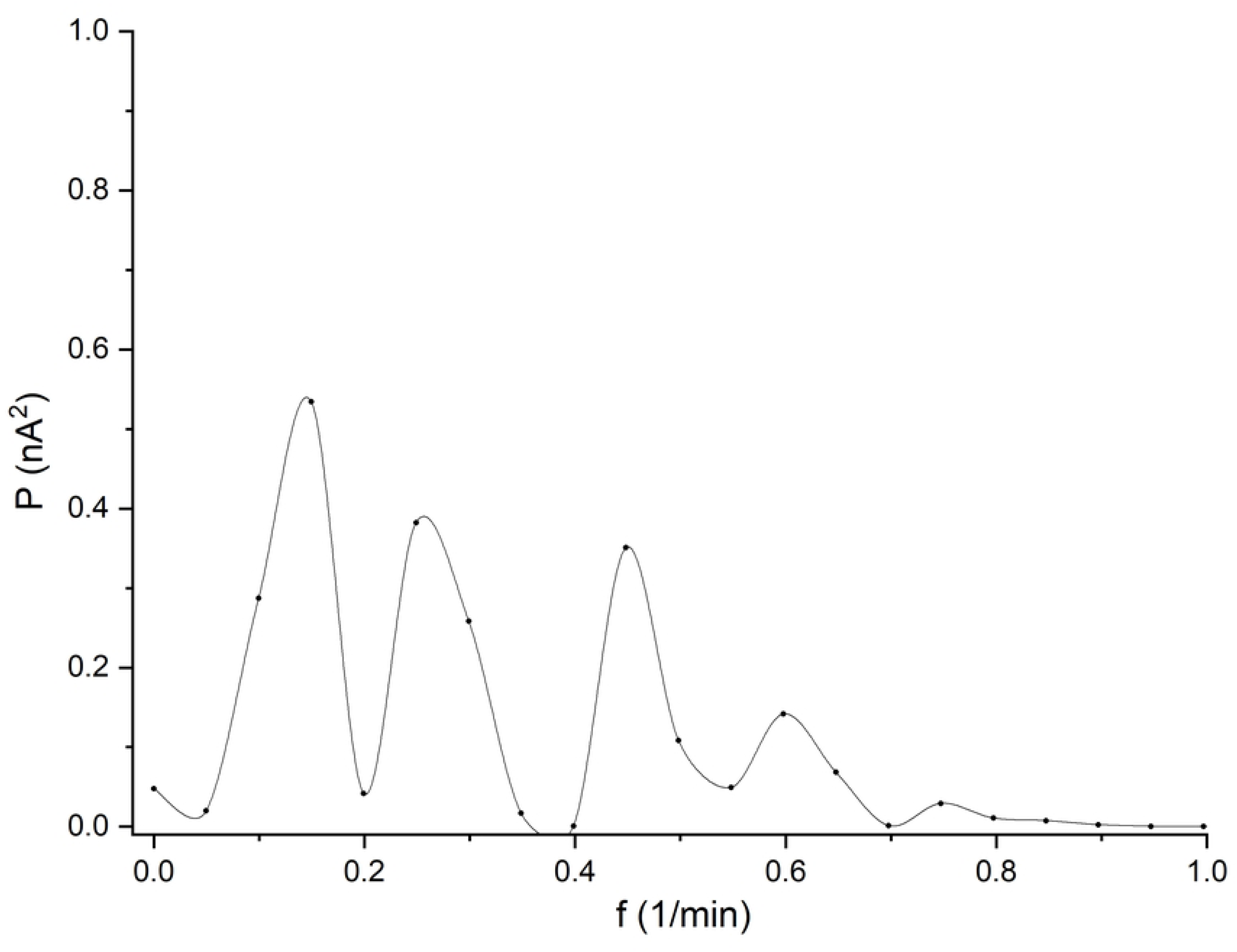

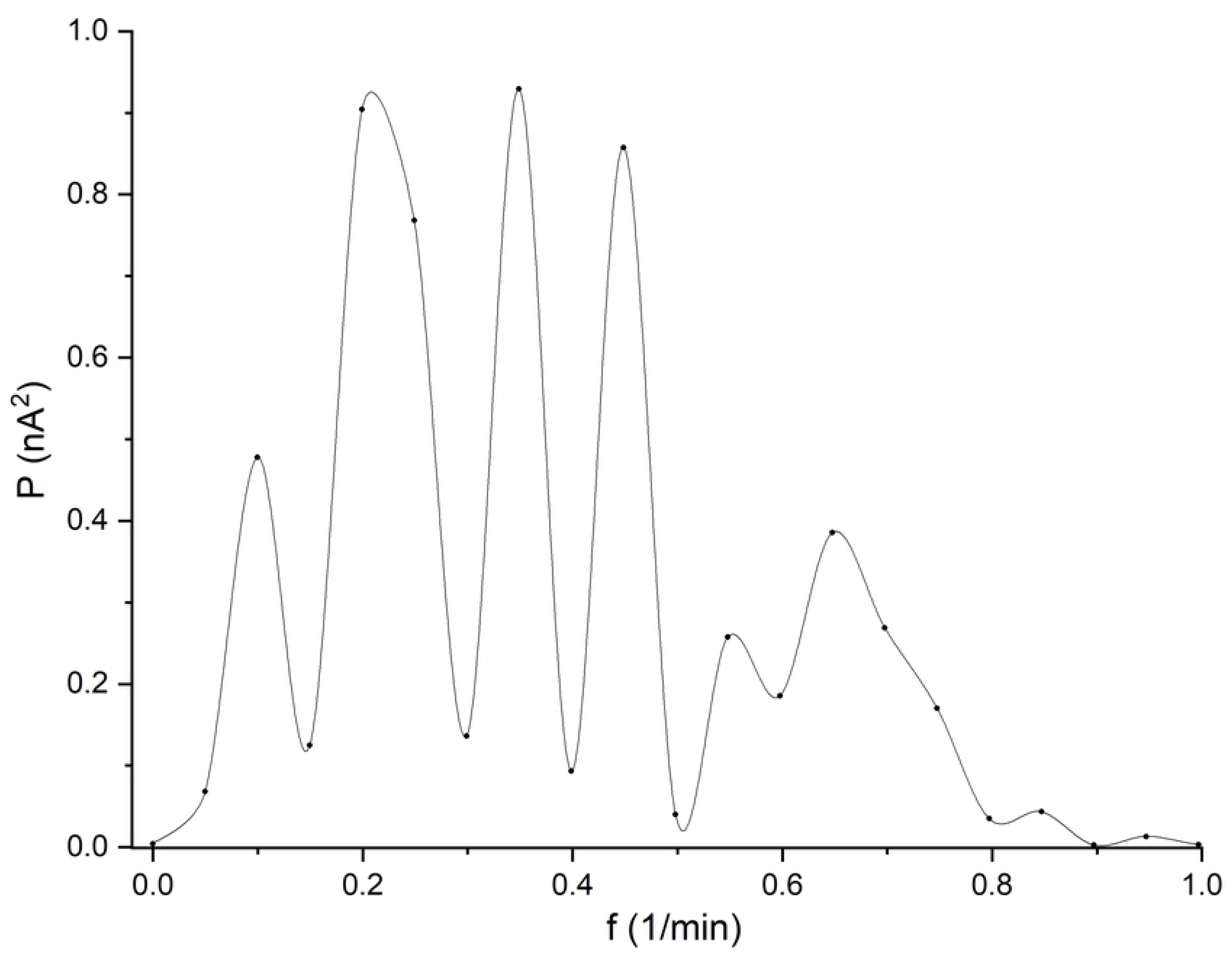

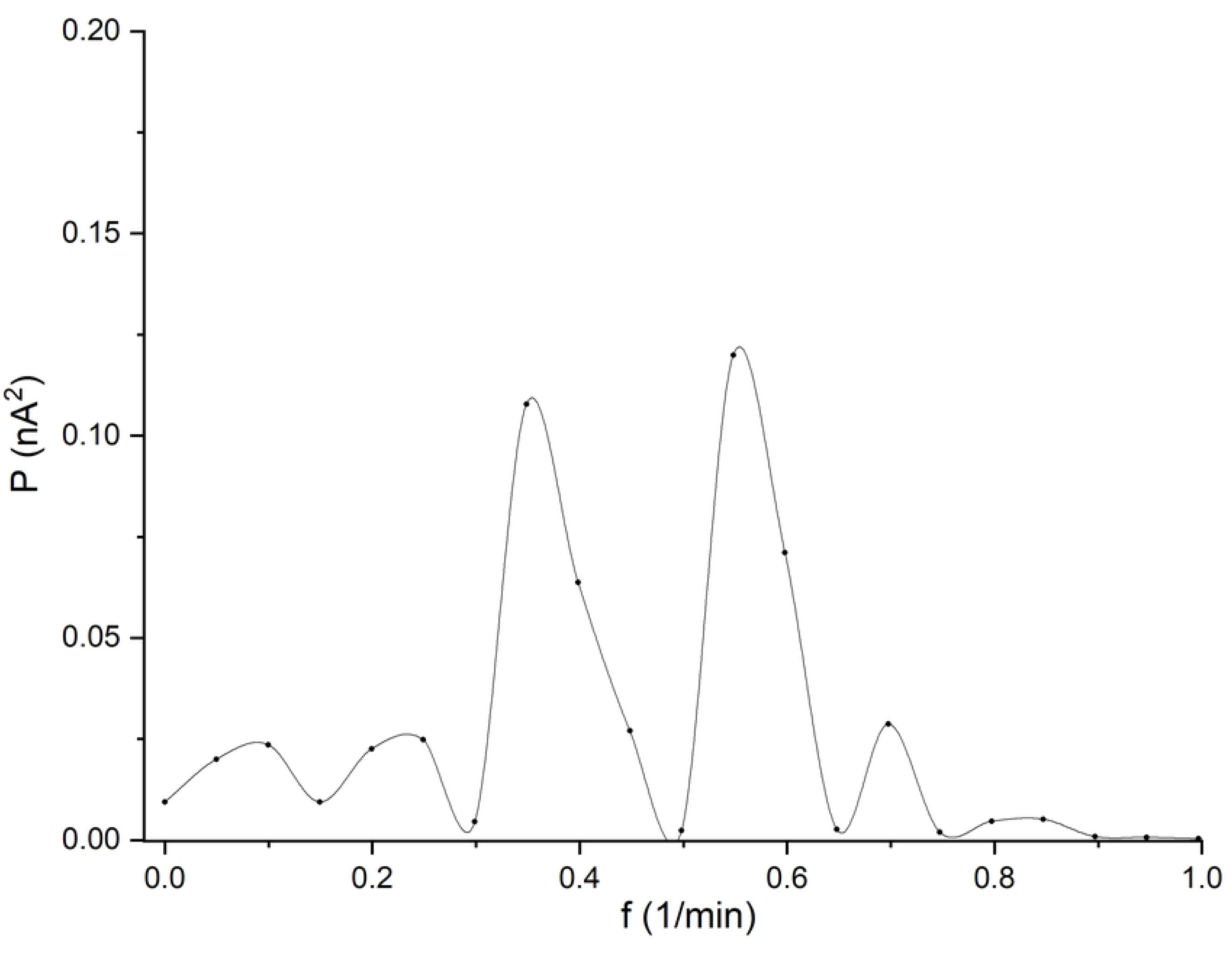

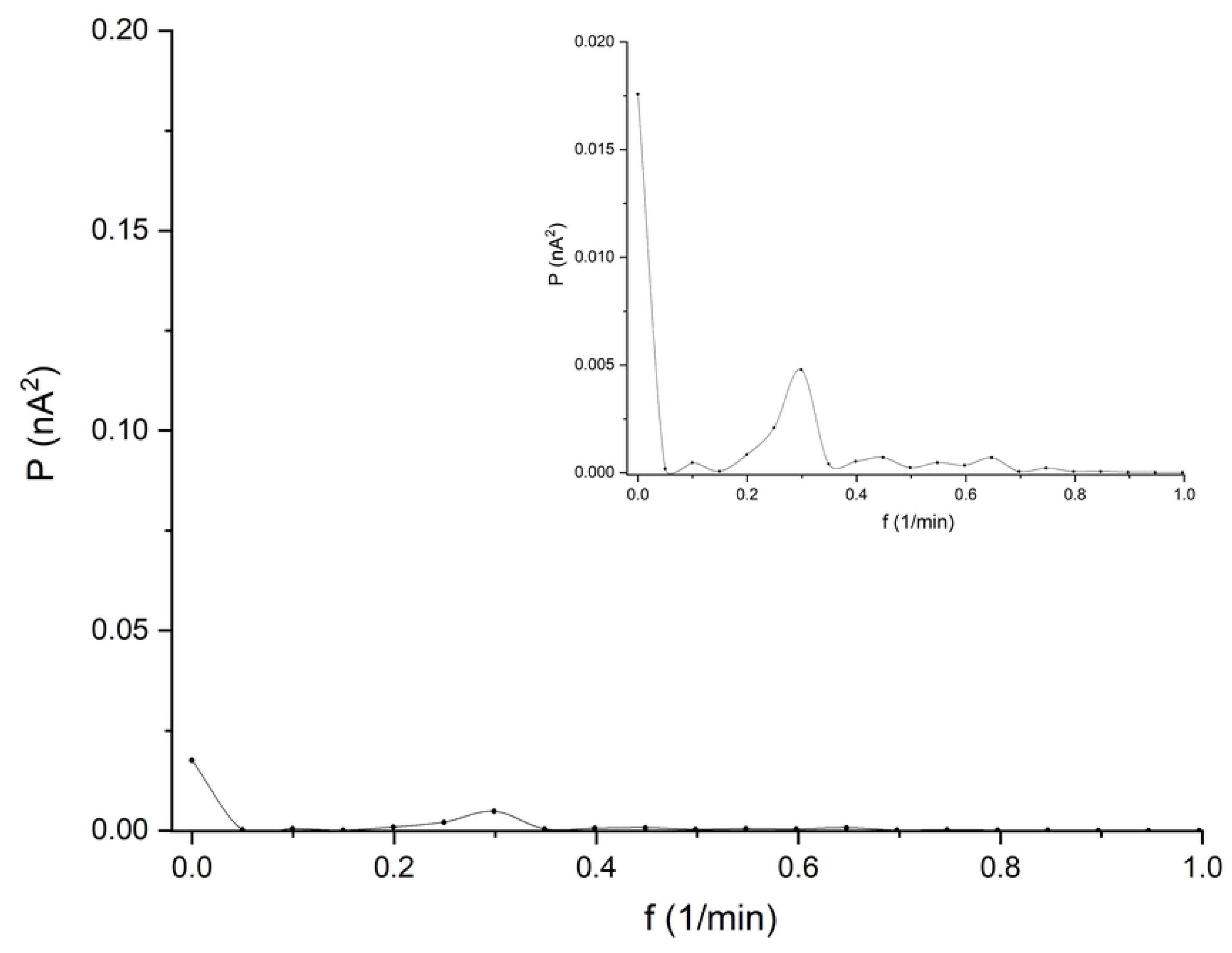

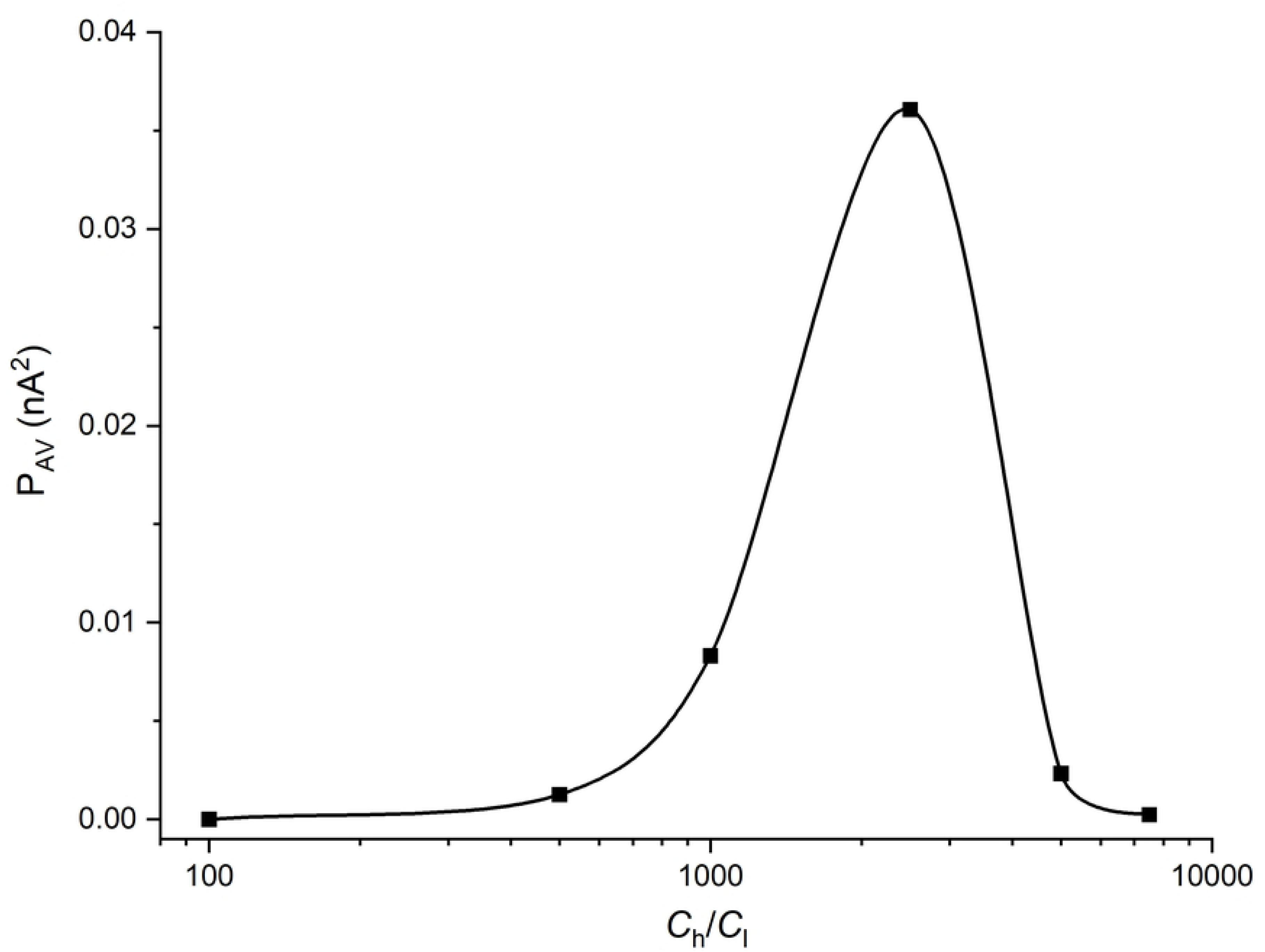

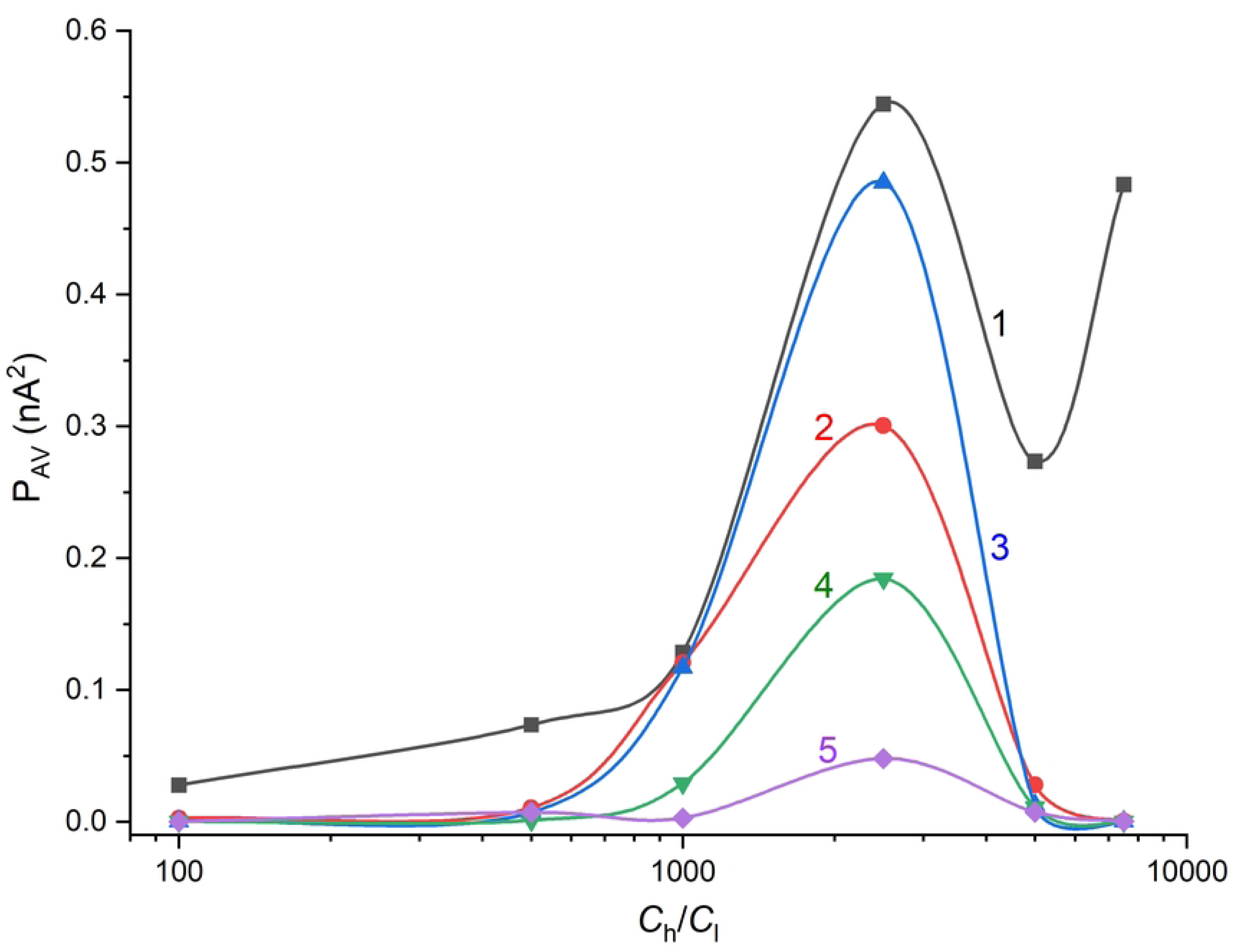

